# Best Practices for Pre-Processing Large-Scale ATAC-Seq Data: Why Epidemiological Studies Demand Greater Stringency

**DOI:** 10.64898/2026.10.22.760939

**Authors:** Philippa M. Wells, Marina Flores-Payan, Barry Chioza, Joe Burrage, Ann C. Babtie, Sarah J. Marzi, Leonard C. Schalkwyk, YouthGEMS Consortium, Emma Dempster, Jonathan Mill, Eilis Hannon

## Abstract

Assay for transposase accessible chromatin using sequencing (ATAC-seq) and related techniques provide critical insight into the epigenetic mechanisms that regulate gene expression in health and disease. Accurate identification of biologically meaningful chromatin accessibility signals remains an underexplored analytical challenge. Advances in sequencing technologies have increased both the magnitude and complexity of ATAC-seq datasets, leading to demand for scalable, reproducible and robust computational analysis frameworks. Peak calling, which identifies regions of chromatin accessibility, is a critical step that underpins all downstream quantitative analyses. However, there is a lack of consensus regarding the optimal peak calling strategy. Here, we present a novel bioinformatics pipeline PIPPA (Pipeline for Integrative Peak calling in Population-scale ATAC-seq), for quality control and peak set curation of large-scale ATAC-seq datasets. Using two independent cell-type sorted ATAC-seq datasets, we show that peak calling performance is dataset-dependent and that no single significance threshold performs optimally across all scenarios. We provide practical recommendations for parameters to generate high-confidence chromatin accessibility profiles. Our peak calling approach is applicable for other genome wide epigenomic assays such as ChIP-seq and CUTCTag. By standardising this process, the PIPPA pipeline enhances epidemiological studies focused in identifying epigenetic dysregulation associated with development and disease.

## Introduction

Epigenetic regulation, including chromatin structure, plays a central role in controlling gene expression and cellular identity by modulating the accessibility of genomic DNA to regulatory elements such as transcription factors.^1^ As regulatory dysfunction is known to play a role in many disease phenotypes^2,3^, mapping chromatin accessibility in the appropriate cell type is essential for identifying the molecular mechanisms underpinning the disease.

The assay for transposase accessible chromatin using sequencing (ATAC-seq) is a high-throughput method that enables unbiased, genome-wide profiling of chromatin accessibility.^4^ Unlike approaches such as chromatin immunoprecipitation sequencing (ChIP-seq)^5^ or Cleavage Under Targets C Tagmentation (CUTCTag)^6^, which target a specific epigenetic modification, ATAC-seq provides a broad overview of the gene-regulatory activity landscape across the genome and its relative simplicity, high sensitivity, and low input requirements have made it a widely adopted technique. The assay works by using a Tn5 transposase to insert sequencing adapters into regions of accessible chromatin enabling the identification of open chromatin regions as genomic regions statistically enriched for sequencing reads, commonly referred to as “peaks”.

Since its introduction, ATAC-seq has been widely used to generate maps of chromatin accessibility across diverse tissues, cell types and experimental conditions. Early studies characterised regulatory landscapes in relatively small cohorts, providing fundamental insights into cell type-specific gene regulation and epigenome organisation.^7,8^ However, as with other genomic data, the use of ATAC-seq in epidemiological studies to identify dysregulation associated with disease requires large datasets to sensitively detect inter-individual variation and overcome the multiple-testing burden inherent in genome-wide analyses. As the scale and complexity of ATAC-seq datasets have increased, the field has progressed beyond descriptive profiling towards identifying chromatin accessibility differences associated with a broad range of biological processes and diseases.^9,10^ Cell type-specific epigenomic approaches using purified populations of nuclei represent an important future direction providing cell type-resolved profiles and overcoming the cellular heterogeneity that can obscure regulatory signals in bulk tissue^11–13^, and limited coverage of single cell studies.

The rapid growth of chromatin accessibility datasets has not been accompanied by the development of bioinformatics tools specifically tailored for their analysis in large epidemiological datasets.^14^ Many studies have relied on methods originally developed for related assays, such as ChIP-seq and DNase I hypersensitive sites sequencing (DNase-seq), despite fundamental differences between ATAC-seq and other chromatin profiling techniques.^15,16^ The ENCODE ATAC-seq pipeline is widely regarded as the preferred computational workflow for the analysis of chromatin accessibility datasets.^17^ However, whilst is has been considered a gold standard it was developed at a time when the sample type, size and scale of available datasets were considerably smaller than those generated today. As dataset sizes have increased, workflows originally developed for smaller datasets may not scale efficiently for large datasets, or more general pipelines omit important quality control (QC) steps specific to ATAC-seq. Thus, there remains a need for a comprehensive, end-to-end workflow capable of reliably processing large scale ATAC-seq datasets from raw sequencing reads to quantitative chromatin accessibility profiles in a reproducible manner.

Within this framework, peak calling is a critical step in ATAC-seq data analysis, identifying the genomic regions corresponding to open chromatin signal and providing the basis for downstream analyses, including differential accessibility between conditions, transcription factor motif enrichment analysis and inference of gene regulatory networks. Typically, peaks are called directly from the dataset being analysed using statistical approaches that distinguish enriched accessibility signals from technical variation and background noise. Among these methods, Model-based Analysis for ChIP-Seq (MACS)^18^, originally developed for ChIP-seq data, remains one of the most widely used peak calling methods in ATAC-seq analyses. Despite its widespread adoption, there is limited empirical data or consensus on how MACS should be optimally applied to ATAC-seq and other genome wide epigenomic data, especially as studies increase in scale.

Here we present a novel ATAC-seq preprocessing and peak calling pipeline, PIPPA (Pipeline for Integrative Peak calling in Population-scale ATAC-seq), alongside guidelines for applying MACS peak calling in large-scale datasets. To validate the pipeline and assess the robustness of the resulting peak sets, we leveraged two large, independent cell type specific ATAC-seq datasets generated from post-mortem prefrontal cortex (PFC) tissue using fluorescence activated nuclei sorting (FANS)^12^, enabling a systematic evaluation of MACS peak calling performance across different biological contexts and dataset sizes. We found that the standard Q value settings led to inflated numbers of detected peaks, an effect which is exacerbated as sample size increases. Based on these observations, we propose a generalizable framework for generating reproducible ATAC-seq peak sets from large-scale data by empirically deriving a more robust significance threshold.

## Methods

### Summary of cohorts

Our study comprised nuclei-sorted ATAC-seq data generated from postmortem pre-frontal cortex tissue in two independent studies referred to here as Dataset 1 and Dataset 2 (**Supplementary Table 1**), from the Brains for Dementia Research cohort^19^ and Autism BrainNet^20^, respectively. Dataset 1 included samples from 161 individuals, whilst Dataset 2 included samples from 49 individuals. Dataset 2 is comprised of QC passing samples from our study of DNA methylation in autism using our BrainFANS DNA methylation pipeline.^21–23^ Human cortex tissue was collected under ethical approval and transferred to our care through Materials Transfer Agreements. The study was conducted in concordance with the ethical standards of the Declaration of Helsinki.

### Isolation of neural nuclei from post-mortem brain tissue

To isolate neural nuclei from post-mortem human prefrontal cortex (PFC) samples we performed FANS following an optimised protocol developed by our group.^24^ To isolate neural nuclei from post-mortem human prefrontal cortex (PFC) samples we performed FANS following an optimised protocol developed by our group.^24^ To isolate up to four populations of purified cells, a series of antibodies were used including NeuN-Alexa488, anti-SOX10 NL577-conjugated, and/or anti-IRF8 APC-conjugated antibodies. Briefly, our gating strategy firstly applied NeuN to gate neuronal nuclei (NeuN+), followed by SOX10 to gate oligodendrocytes (NeuN-/SOX10+) and then IFR8 to gate microglia (NeuN-/SOX10-/IRF8+), with the triple negative population (NeuN-/SOX10-/IRF8-), which is enriched for astrocytes, also collected.

### Chromatin accessibility profiling (ATAC-seq)

Chromatin accessibility was profiled using ATAC-seq (Assay for Transposase-Accessible Chromatin sequencing) to identify open chromatin regions across the genome. As described in Chioza *et al* 2026^12^, approximately 100,000 nuclei were pelleted (1,000 × g, 5 min, 4 °C), resuspended in BAMBANKER™ Cryopreservative, and frozen at −20 °C. For library preparation, nuclei were thawed, washed with PBS, and processed following the Omni-ATAC protocol (28), including transposition and PCR amplification using NEBNext 2X Master Mix and custom Nextera ATAC-seq primers. Libraries were purified using double-sided SPRI bead selection, quantified using the NEBNext Library Quant Kit, pooled equimolarly, and sequenced on an Illumina Novaseq X at 50bp paired end by the Exeter Sequencing Service.

#### ATAC-seq data preprocessing

Our bespoke ATAC-seq pipeline, PIPPA, was developed following the same framework as our established DNA methylation pipeline.^21^ This is structured as three quality control stages that aim to confirm i) the quality of the data, ii) the sample identity and iii) the assigned cell type, respectively (**Supplementary Figure 1**).

##### Sample quality control

Raw reads provided as fastq files were processed as follows. First, sequencing summary metrics were extracted using FastQC^14^ (version 0.12.1). Sequencing adapters (using a prespecified custom list and *–adapter* fasta flag) and low-quality bases were trimmed from each end of the read using Fastp (version 0.23.4) up to a specific maximum length (using flags *-b* and *-B*). Trimmed reads were summarised with FastQC and aligned to the human reference genome (NCBI GRCh38) using Bowtie2^16^ (version 2.5.4). Aligned reads were converted to SAM format, sorted and indexed using SAMtools^17^ (version 1.21), and reads that were unmapped, had a secondary alignment or failed the minimum quality score (MAPQ score <30) were excluded. Duplicated reads were marked and removed using SAMtools and Picard^18^ (version 3.3.0) *MarkDuplicates*. The output for each sample was a BAM file containing non-duplicated, mapped in proper pairs, filtered, aligned reads. To assess the quality of the sequencing data, the total number of read pairs and the number of distinct read pairs were counted using SAMtools and SAMStat^19^ (version 0.2.2).

A series of additional metrics were calculated to assess the quality of the ATAC-seq libraries. Three ENCODE library complexity metrics (non-redundant fraction, PCR bottlenecking coefficient 1 and 2) were calculated as described on the ENCODE website^10^. Non-redundant fraction (NRF) measures the proportion of distinct uniquely mapped reads among all reads. PCR bottlenecking coefficient 1 (PBC1) quantifies the proportion of uniquely mapped genomic locations which contain exactly one read, while PBC2 compares the number of locations with exactly one uniquely mapped read to those with exactly two uniquely mapped reads. The fragment size distribution was characterised to confirm decreasing periodicity by classifying aligned reads as those capturing nucleosome free (Nuc-Free) and mono (Mono-Nuc), di-, tri- and quad-nucleosomes regions using the R (version 4.3) package ATACseqQC^20^ (version 1.26.0) and converted into proportions that capture the abundance of each nucleosome size. Thresholds for the proportion NFR and MNC were evaluated as part of this work to automate and standardise the review of the fragment distribution. Peak calling was performed on sample-level BAM files with MACS (Model-based Analysis of ChIP-seq; version 3.0.2) as follows: *macs3 callpeak –f BAMPE –nomodel –keep-dup all –g 2.9e9 -q 0.05*. Peaks overlapping ENCODE hg38 blacklisted regions (version 2)^21^ were excluded.

BAM files containing filtered aligned reads (with duplicates retained) were processed using deeptools^22^ (version 3.5.5) *bamCoverage* and *bigWigCompare* to generate bigWig files. These were used to calculate Transcription start site (TSS) enrichment scores as the ratio of aggregated read density centred on TSSs relative to flanking background regions. TSS regions were obtained from refSeq hg38 database.^33^ Quality metrics were collected and summarised in an R Markdown HTML report. Thresholds used to classify samples as passing or failing each QC metric with associated counts of failed sample for each step are specified in **Supplementary Table 1**.

##### Sample identity check

The second stage of the pipeline assesses sample identity using sex and genotype data, where available. To verify labelled sex, aligned bam files were processed as follows. First, aligned reads to the sex chromosomes were extracted into separate BAM files. Peaks were then called on these using MACS^34^: *macs3 callpeak –f BAMPE –nomodel – keep-dup all –g 2.9e9 -q 0.0001*. Bedtools^5^ (version 2.31.1) *closest* was used to keep the nearest upstream peak that did not overlap with the gene body of X chromosome genes XIST and FIRRE. Bedtools *intersect* was used to exclude peaks called on chromosome Y that overlap pseudoautosomal regions. These are defined as those regions where chromosome X and Y share homologous sequences^23^. For each sample, the number of reads that overlap X and Y chromosome peaks were counted using Bedtools *intersect.* Samples were assigned a predicted sex based on the level of signal on the Y chromosome relative to the signal in XIST and FIRRE annotated peaks, with higher signal on the Y chromosome indicating “male/M” and lower signal indicating “female/F”. To verify sample identity using matched imputed genotype data available, BAM files containing filtered aligned reads (with duplicates retained) were processed using GATK (version 4.6.1.0) *BaseRecalibrator* and *ApplyBQSR*^25^ to generate recalibrated base quality reads. Base calls were compared against the genotype data available for each sample using VerifyBamID^22^ (version 1.1). Output metrics FREEMIX and CHIPMIX were used to assess potential sample contamination and sample swaps, respectively. A threshold of 0.1 was applied to both metrics, indicating that ≥10% of reads at reference sites were inconsistent with the expected genotype. The results of these sample-identity checks were summarised in a second R Markdown HTML report.

##### Cell type identity check

To confirm that the cell type listed for the samples’ metadata matched the experimental profile, peaks were called on merged samples separately for each cell fraction. Merged, sorted BAM files were generated with SAMtools, and peaks were called on the 4 cell types-specific large BAM files using MACS as follows: *macs3 callpeak –f BAMPE – nomodel –g 2.9e9 –keep-dup all -q 0.05.* Peaks overlapping ENCODE hg38 blacklisted regions (version 2) were excluded. Cross cell type consensus peaks were generated using GenomicRanges R package (version 1.54.1), where peaks within 150 bp distance were considered to be the same peak and merged. A per-sample count matrix of reads falling within the cross cell type consensus peak set was generated using FeatureCounts and RSubRead^25,26^ and was used as input to edgeR R package^27^ (version 4.0.16). EdgeR was used to filter low-counts peaks with less than 2 counts per million in fewer than 2 samples and to perform differential analysis of cell type. Principal component analysis (PCA) was performed on normalised counts from the top 1000 differential peaks across cell types (ranked by p-value) to align the major axes of variation with cell type differences. The silhouette coefficient was calculated using the R package cluster^28^ (version 2.1.8.2) and the first 5 PCs to quantify the similarity of each sample to their cell type group identify potential swaps or unsuccessful FANS isolation experiments. Silhouette analysis measures how similar a data point is to others within its cluster.^41^ In our application, clusters were defined by sample cell type labels. The silhouette score ranges from 1 to -1, where a score near 1 indicates that the sample is closely aligned to its assigned cluster, while score near -1 indicates that the sample aligns more closely with a different cluster (i.e. a cell type other than what it is labelled as). The results of these analyses, alongside the final QC status of each sample (pass or fail) were summarised in a third R Markdown HTML report.

### Peak calling

#### Peak calling across increasing sequencing depths

To profile the impact on peak calling of increasing sequencing depth, down-sampled BAM files were generated across a range of read depths. For a chosen cell type group, the BAM files for all samples that passed all three stages of the QC pipeline were merged and sorted using SAMtools *merge* and *sort*, respectively. Then, the merged BAM file was down-sampled to produce BED files containing between 100 million and 1 billion read pairs at 100 million increments, using MACS *randsample* with parameters *–f BAMPE --seed 2*. Peak calling was performed on these using MACS: *macs3 callpeak* -*f BEDPE* –*nomodel* –*g 2*.*SeS* –*keep*-*dup all* at Q value thresholds 0.05, 0.01 and 0.00001.

#### Genomic characterisation of peaks

Overlap between peak sets was quantified using the R GenomicRanges package. Genomic element annotation was performed using the R ChIPseeker package^42^ (version 1.38.0), with default parameters and TxDb.Hsapiens.UCSC.hg38.knownGene (version 3.18.0) and org.Hs.eg.db (version 3.18.0) databases. Highly expressed brain genes were obtained from the Lieber Institute for Brain Development ^43^ study of the transcriptome in Schizophrenia, lifespan prefrontal cortex samples.^44^

#### Identification of the elbow of the peak number versus Q value curve

Peaks were called on the merged BAM file including samples that passed QC for each cell fraction at Q<0.05. Peak numbers versus Q value were plotted from Q < 0.05 to Q < 1e^-100^. For this threshold discovery exercise, peak Q values were subset from the larger Q<0.05 peak set. Segmented analysis to mathematically identify the elbow of the curve was performed using R package segmented^34^ (version 2.2.1) using peak threshold increments of 2 1e^-n^ Q values (e.g. 1e^-5^, 1e^-7^, 1e^-9^, …).

#### Association between sequencing depth and elbow of the peak number versus Q value curve

Association between the curated Q value for each peak set (derived as described above from the elbow of the peak number versus Q value curve, indicated using segmented analysis), and the sequencing depth (total aligned fragments) of the merged bam file from which each peak set was called was assessed for all four cell fractions across both datasets, and separately. A linear model was applied without covariates using the formula *mod1 <-lm*(*Q ∼total*_*aligned*_*fragments*, *data*=*depth*), in the lme4^46^ R package (version 2.0-1).

#### Visualisations

Figures were created using ggplot2 R package^47^ (version 4.0.3), and multipaneled figures were collated in R using the cowplot package (version 1.2.0)^48^.

#### Pipeline technical specifications & availability

The pipeline was designed for execution on an HPC system using the SLURM job scheduler. It comprises a series of main Bash scripts that can be executed from the command line, with individual processing steps implemented through additional Bash, Python or R scripts. File paths, installed software, reference files and QC metrics thresholds are specified in configuration files (*config*.*txt* and *config*.*r*). Unless otherwise stated, software dependencies required for the pipeline are managed using a conda environment that can be directly installed from a requirements file provided with the pipeline. The pipeline can be accessed from our GitHub repository (https://github.com/ejh243/BrainFANS).^22^

## Results

### Novel ATAC-seq QC framework provides orthogonal assessment of sample and cell type identity

We developed a three-stage preprocessing and quality control pipeline for cell-sorted ATAC-seq data (**Supplementary Figure 1**), PIPPA, building on the previously established pipeline for DNA methylation data.^49^ Stage 1 assesses overall sequencing and ATAC-seq assay quality. Stage 2 verifies that sample identity aligns with their corresponding individual label by checking concordance with sex and genotype information. Stage 3 assesses whether the observed chromatin accessibility profile is consistent with the FANS-isolated cell type. By automating the calculation of quality control metrics within each stage of the pipeline, we provide a scalable and reproducible framework for assessing large ATAC-seq datasets (**Supplementary Table 1**). To validate PIPPA, we applied it to two large cell-sorted datasets with data from four isolated cell fractions (see **Methods**).

Stage 1 of PIPPA calculates metrics common to many sequencing pipelines, including the number of aligned reads, alongside established metrics from the ENCODE consortium ATAC-seq pipeline including measures of PCR bottlenecking (PBC1;PBC2) and the non-redundant fraction (NRF) (ENCODE Project Consortium, 2012)^17^. Across these three metrics, both datasets exhibited unimodal distributions spanning the two highest-quality categories (**Supplementary Figures 2,3**). We therefore set the filtering the thresholds to retain samples falling within either of these categories for subsequent analysis.

A successful ATAC-seq experiment is expected to exhibit periodicity in the fragment size distribution with a prominent nucleosome-free mode (∼50-100 bp), corresponding to fragments originating from accessible chromatin regions, followed by modes of progressively decreasing amplitude representing mono-nucleosome (∼200 bp) and di-nucleosome (∼400 bp) fragments. This characteristic of the data is typically assessed manually, making it subjective and not scalable to large datasets. To automate this process, we developed quantitative metrics that capture the fragment size periodicity by quantifying the proportion of reads within each nucleosome fragment size bin (see **Methods**). Across all the samples, clear periodicity was observed (**Figure 1A**), with progressively smaller proportions of longer fragments. Across samples, there was wide variability in these metrics, indicating that they can capture differences in behaviour at sample level. Based on visual inspection of the fragment distribution for samples at the extremes we suggest minimum thresholds of 0.25 for the proportion of nucleosome-free fragments and 0.2 for the proportion of mono-nucleosome fragments to detect samples lacking the expected nucleosomal periodicity (**Supplementary Figure 4**). Closer inspection of these samples found they typically showed fragment length distributions dominated by a single mode.

**Figure 1.**
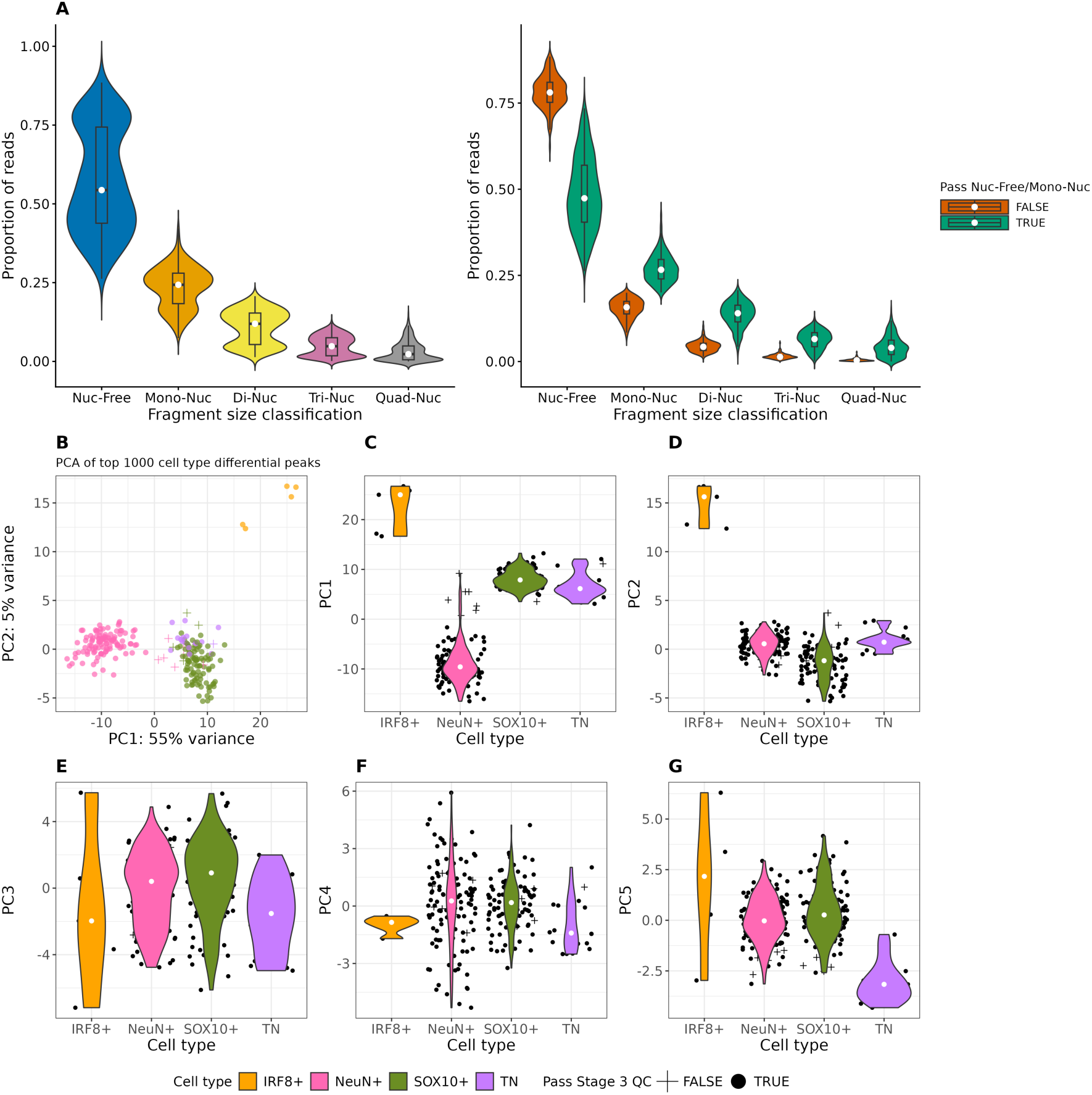
Bespoke quality control metrics confirm successful ATAC-seq experiment in cell-sorted samples. **A**) Classification of fragment sizes overall and classification of fragment sizes split by whether samples pass thresholds for Nuc-Free (0.25) or Mono-Nuc (0.2). Nuc-Free: Nucleosome-free region, Mono-Nuc: mono-nucleosomes, Di-Nuc: di-nucleosomes, Tri-Nuc: tri-nucleosomes, Quad-Nuc: quad-nucleosomes. **B**) Scatterplot of the first two principal components where each point represents a sample, the colour of the point represents the labelled cell type nuclei fraction, and the shape indicates whether the sample is clustering with its labelled cell type fraction based on the silhouette coefficient (I.E passing/failing stage 3 QC). Principal component analysis (PCA) performed on top 1000 cell type differential peaks. **C-G** Violin plots of the first 5 principal components grouped by nuclei fraction. Data are shown for Dataset 1.

Before QC, Dataset 1 included 445 samples across the four cell types, whilst Dataset 2 included 173 samples. Across the two datasets, 201 and 51 samples failed Stage 1 respectively, with the majority failing on a single QC metric (80% and 62% of the failed samples, respectively). In Dataset 1, the proportion of mono-nucleosome fragments was the most frequent cause for sample failure, demonstrating that this metric captures characteristics of ATAC-seq data quality not identified in current pipelines. In dataset 2 the most common cause for failure was the number of aligned reads, highlighting the technical variation in ATAC-seq experiments and justifying the inclusion of multiple QC metrics that assess a range of features. To explore whether poor performance on one metric predicts poor performance on another, we examined the correlations between them (**Supplementary Figure 5**). Across both datasets, the ENCODE QC metrics NRF, PBC1 and PBC2 were strongly positively correlated with each other and showed moderate correlations with aligned read count and fragment distribution metrics. Alignment rate was the most distinct metric, showing at most a weak correlation with the percentage of nucleosome free fragments. This metric is primarily a sequencing-quality metric that confirms reads are correctly mapped to the reference genome, whereas the correlations amongst ENCODE QC metrics reflect a shared dimension of ATAC-seq-specific library quality.

### Novel automated approach for ATAC-seq QC confirms sample and cell type identity

Having confirmed overall ATAC-seq data quality, Stage 2 of the pipeline assessed sample identity using two independent approaches. First, sex was inferred from the data by quantifying the signal in Y-chromosome linked peaks relative to signals at XIST and FIRRE on the X chromosome (see **Methods**). Based on sex chromosome signals, samples cluster into two distinct groups, representing males and females enabling a comparison of inferred and recorded sex. Second, ATAC-seq data was compared to matched genotype data from the same individual using *verifyBamID* to assess genotype concordance and detect potential contamination. Samples showing mismatches with recorded sex and genotype data were excluded (**Supplementary Figure 6**). In Dataset 1, 68 samples failed Stage 2, of which 52 didn’t align with their matched genotype information, 2 had their sex incorrectly predicted and 14 failed both checks.

The final stage (stage 3) of PIPPA is bespoke to the cell type-specific nature of the data and confirms that the isolated fractions represent distinct cell types. Here we assess whether each sorted nuclear fraction exhibited the expected cell type-specific chromatin accessibility profile. Principal component analysis across peaks that differ between cell types clustered samples by their assigned cell fraction (**Figure 1B**). In both datasets, the first PC (explaining 55% and 32% of the variance respectively) separates the NeuN+ samples from the glial cell populations. The second PC (explaining 5% and 15% of the variance, respectively) separates the IRF8+ samples from the others. In Dataset 1 the third and fourth PCs are not associated with cell type and the fifth PC separates the TN fraction. To objectively decide whether a sample fits the profile of its labelled fraction better than the profile of any other cell type, silhouette coefficients were calculated. Samples with negative silhouette scores (indicating that a sample aligns more closely with another cell type cluster than its labelled sample cell type) calculated across PCs 1 to 5 were excluded, as these may be indicative of either an unsuccessful isolation or mislabelled sample. In Dataset 1, 13 samples didn’t cluster with their assigned cell type. In Dataset 2, only 2 samples were identified as mislabelled samples and were excluded from subsequent analyses.

Across all three stages of the QC pipeline, 222 samples from Dataset 1 passed all QC criteria. Samples that passed were associated with higher read depth (mean 1.21 × 10^8^ vs 9.5 × 10^7^; p value < 0.001), a more balanced nucleosome free-to-mono-nucleosome ratio (mean 1.8 vs 4; p value < 0.001) and higher TSS enrichment (mean 2.7 vs 2.2; p value < 0.001) than failed samples (**Supplementary Figure 7**). In Dataset 2, 120 samples across all cell types passed all three stages of the QC pipeline. These samples were also associated with higher read depth (mean 2.91 × 10^8^ vs 1.38 × 10^8^; p value < 0.001).

### Higher sequencing depth leads to inflated numbers of peaks

Read depth is an important consideration in sequencing study design, conferring power for peak discovery and downstream statistical analysis. In principle, increasing depth should identify additional peaks, with the rate of new peaks eventually decreasing as peak detection reaches completion. Leveraging our largest dataset and focusing the NeuN+ nuclei fraction (n= 109 samples, mean read depth = 1.22 × 10^8^), we investigated the relationship between the number of peaks and sequencing depth at a limited range of standard Q value thresholds (0.05, 0.01, 0.00001) by subsampling the reads merged across all samples. Whilst we intended that these analyses would give some insight into the completeness of the peak set, they did not demonstrate evidence of the anticipated plateauing behaviour (**Figure 2A**). Instead, the number of peaks continued to increase, reaching a maximum of 500,000 peaks at the highest sequencing depth (10 × 10^8^) when using Q< 0.05. In total 4.2 × 10^8^ bp of sequence was included in these peaks, capturing 16% of the human genome.

**Figure 2:**
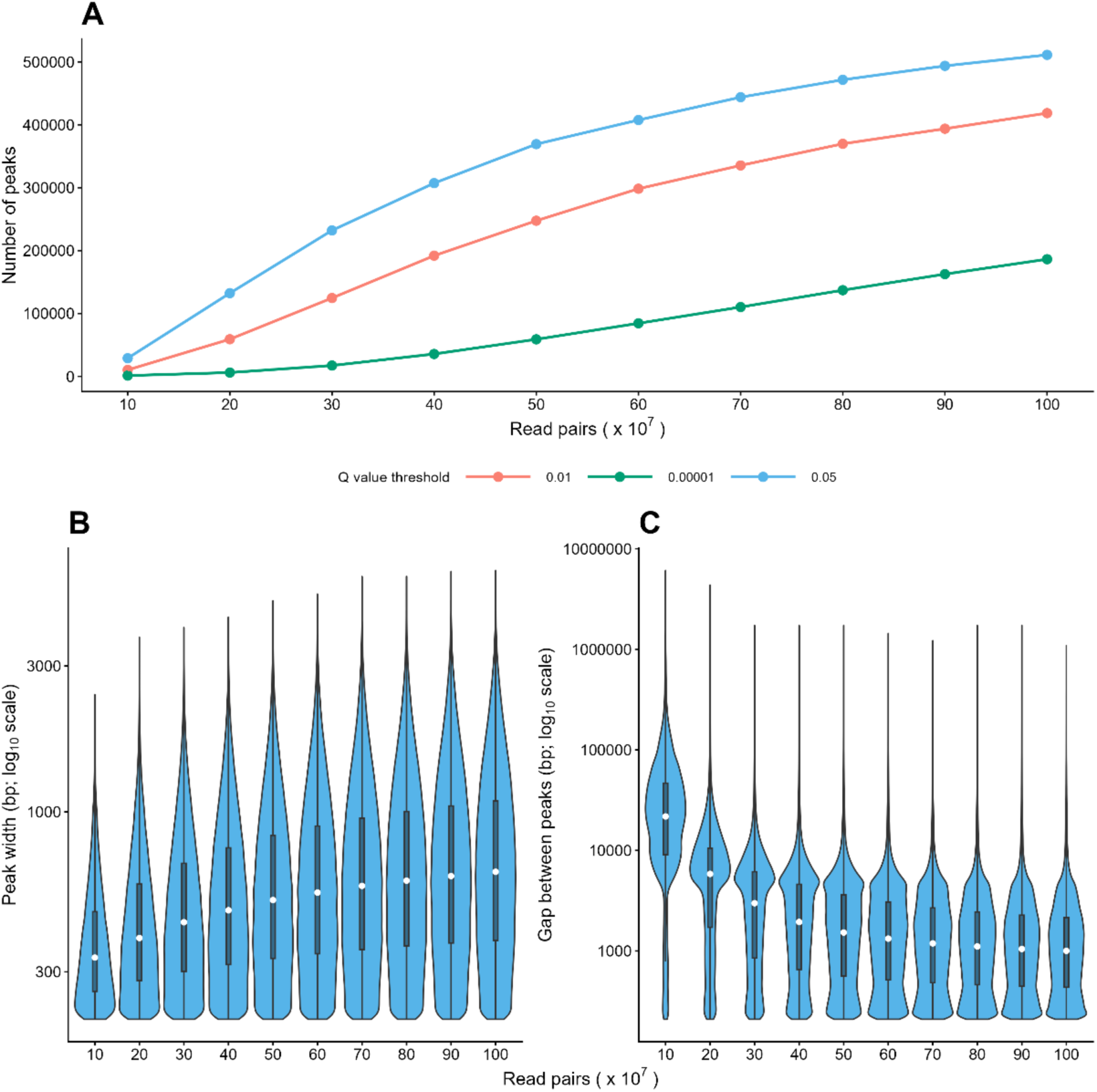
Peak calling exhibits sequencing depth–dependent changes in peak number and characteristics. **A**) Line graphs showing the number of peaks (y-axis) identified across increasing sequencing depths (number of read pairs; x-axis), with each line representing a different Q value threshold. Violin plots showing the distribution of **B**) peak widths (base-pairs; bp) and **C**) inter-peak distance (bp) for peaks called at increasing sequencing depth. Peaks were called using Dataset 1 NeuN samples only; For B) & C), peaks were called at Q < 0.05 threshold.

This behaviour indicates signal saturation, where the read coverage increases across both enriched and background regions, negatively affecting the ability of the algorithm to distinguish true signal from background, leading to false positive peaks. Furthermore, the characteristics of the peak set changed as more sequencing data was included, with peaks becoming wider and denser (**Figure 2B-C**), consistent with peak boundaries becoming indistinguishable from background regions. Splitting the peaks based on size, showed that the inflated number was driven particularly by an abundance of narrow peaks (< 400bp; **Supplementary Figure 8**). The number of peaks was reduced and consequently the degree of inflation minimised for wider peaks compared to narrower peaks.

The continued expansion of the peak set raises concerns about biological interpretability. We therefore examined the relationship between sequencing depth and gene annotation, focusing first on protein-coding genes. In contrast to the total peak number, the number of these genes associated with at least one peak showed evidence of plateauing with increasing sequencing depths (**Supplementary Figure 9**). For example, at default Q value (Q < 0.05), 12,425 protein-coding genes were annotated to at least one peak in Dataset 1 NeuN with 50 × 10^7^ read pairs, nearly doubling the number observed at 10 × 10^7^ read pairs (6,660). This number remained relatively stable at higher sequencing depth, with 13,303 protein-coding genes annotated with 10 × 10^8^ read pairs. The number of peaks associated with each gene increased dramatically with sequencing depth, rising from a mean of 3 peaks per gene at 10 × 10^7^ read pairs, to 17 at 50 × 10^7^, and 23 peaks per gene at 10 × 10^8^read pairs at Q<0.05. In contrast, the number of promoter annotated peaks remained stable. For Dataset 1 NeuN at Q < 0.05, fewer than three promoter-peaks were annotated per gene on average with 50 × 10^7^ read pairs, increasing to just over three at the highest sequencing depth.

We also observed a marked increase in the number of narrow peaks (< 400bp) per gene with increasing sequencing depth. However, this relationship began to plateau once sequencing depth reached 50 × 10^7^ read pairs with a mean of 6 narrow peaks per gene at the default Q value threshold (Q<0.05; **Supplementary Figure 9**). The majority of peaks were associated with genes expressed in the brain, with this proportion remaining consistently high at approximately 70% across sequencing depths. The proportion was greatest at the lowest sequencing depth and showed little change as additional sequencing data were incorporated. Similar findings were observed when not only protein-coding genes were considered (**Supplementary Figure 10**).

To further assess the robustness of peak sets, we evaluated how the peaks overlapped those called in each individual sample. As more sequencing data is included, the sample overlap decreases (**Supplementary Figure 11**). Increasing the amount of sequencing data from 50 × 10^7^ to 10 × 10^8^ read pairs, the mean overlap decreases from 9 to 7 samples (Dataset 1 NeuN Q < 0.05). We note that this varies by peak size, with wider peaks overlapping with more samples on average than narrower peaks (**Supplementary Figure 12**). Furthermore, at higher sequencing depths the proportion of total peaks overlapping one or more samples also increased. Of note, most of these overlaps are found in samples with the most data and therefore the most peaks (**Supplementary Figure 12**). Together, these findings indicate that increasing sequencing depth does not lead to a more complete peak set but instead leads to an inflated set of peaks, altering the composition and genomic characteristics of the OCRs that serve as the basis for downstream analysis.

### More conservative significance thresholds are required for deeper datasets

Consistent with the elevated background signal, we hypothesised that a substantial proportion of peaks represent chance findings. Under this framework, the observed Q value distribution is consistent with a mixture of two components: a true enrichment signal and a false positive component arising from random variation. We therefore reasoned that applying increasingly stringent significance thresholds would preferentially eliminate background peaks, enriching the remaining peak set for genuinely accessible regions and reducing the influence of false discoveries.

To evaluate this hypothesis, we investigated the relationship between Q value and number of peaks detected, up to Q < 1e^-100^, separately in neurons and oligodendrocytes in both datasets (NeuN+ and SOX10+). Across all conditions, the number of peaks initially decreased sharply as the threshold became more stringent, before reaching a plateau (**Figure 3A**), with the precise shape of this relationship varying across datasets, likely reflecting differences in sequencing depth (and hence saturation) and signal-to-noise ratio.

**Figure 3:**
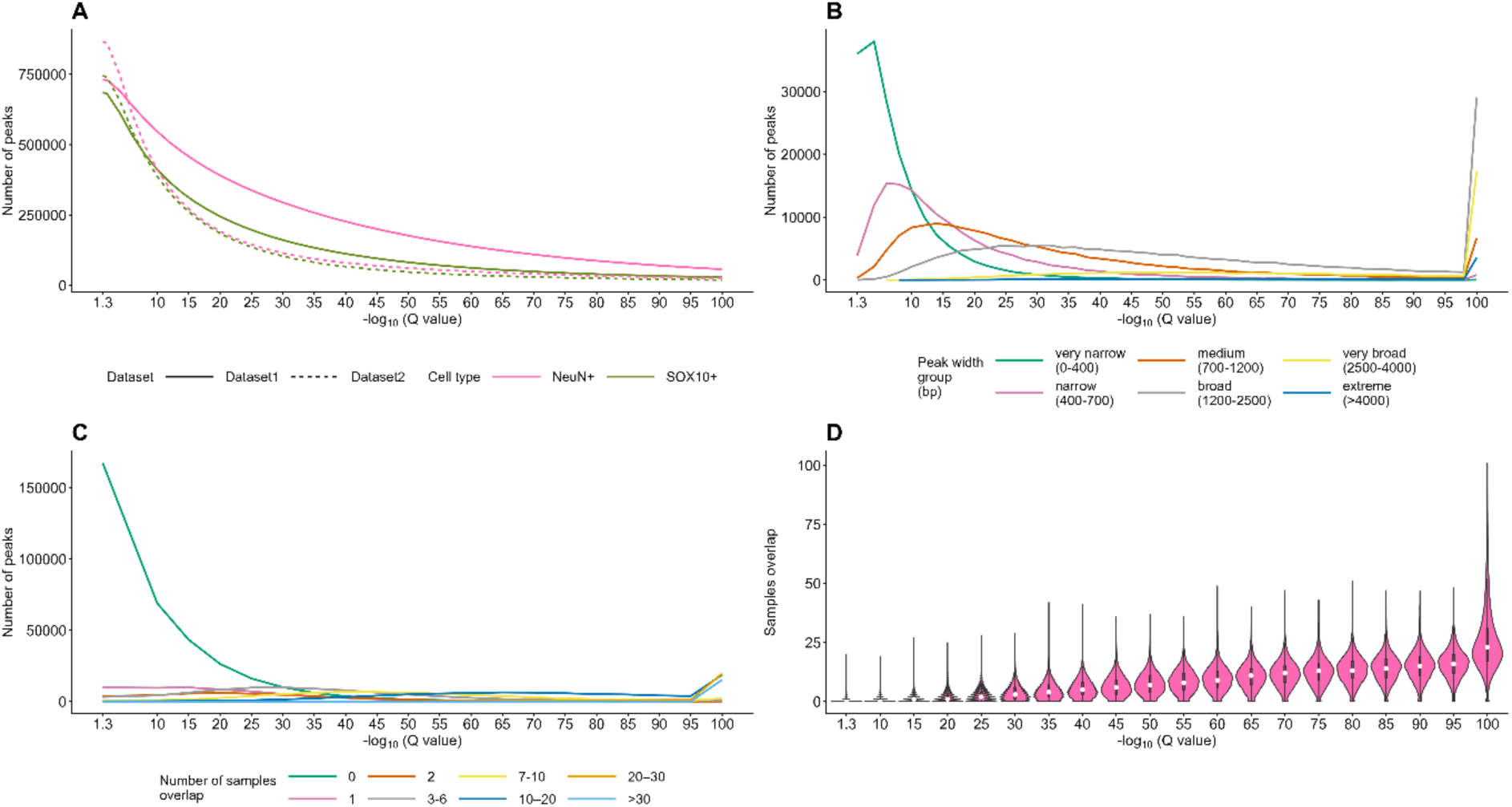
More stringent Q value thresholds alter the characteristics of detected peaks. **A**) Line graph showing the number of peaks (y-axis) at increasingly stringent thresholds (Q value; −*log*_10_; x-axis). Lines represent the different datasets and cell types. **B**) Line graph showing the distribution of peaks grouped by peak width across increasing Q value thresholds. **C**) Line graph showing the distribution of peaks grouped according to the number of individual samples in which they are detected across increasing Q value thresholds. **D**) Violin plots showing the distribution of the number of samples overlapping each peak for peak sets generated at increasing Q value thresholds (x-axis). Panels B, C and D show data for Dataset 1 and NeuN+ cell type only.

Consistent with this interpretation, peaks retained at more stringent thresholds showed greater reproducibility against individual-sample level peaks and a reduced incidence of very narrow peaks (**Figure 3B;3C** and **Supplementary Figure 13**). The distance between neighbouring peaks increased with increasing stringency, consistent with the removal of closely spaced low-confidence peaks. While the number of genes annotated to at least one peak decreased approximately linearly as the Q value threshold became more stringent, the number of peaks per gene decreased more rapidly (**Figure 4A;4B**). The genomic distribution of peaks also changed with increasing stringency, most notably with a greater proportion of the retained peaks annotated as promoters (**Figure 4C**). This suggests that increasing stringency primarily removes additional weaker confidence peaks, while retaining the most strongly associated peaks associated with each gene.

**Figure 4:**
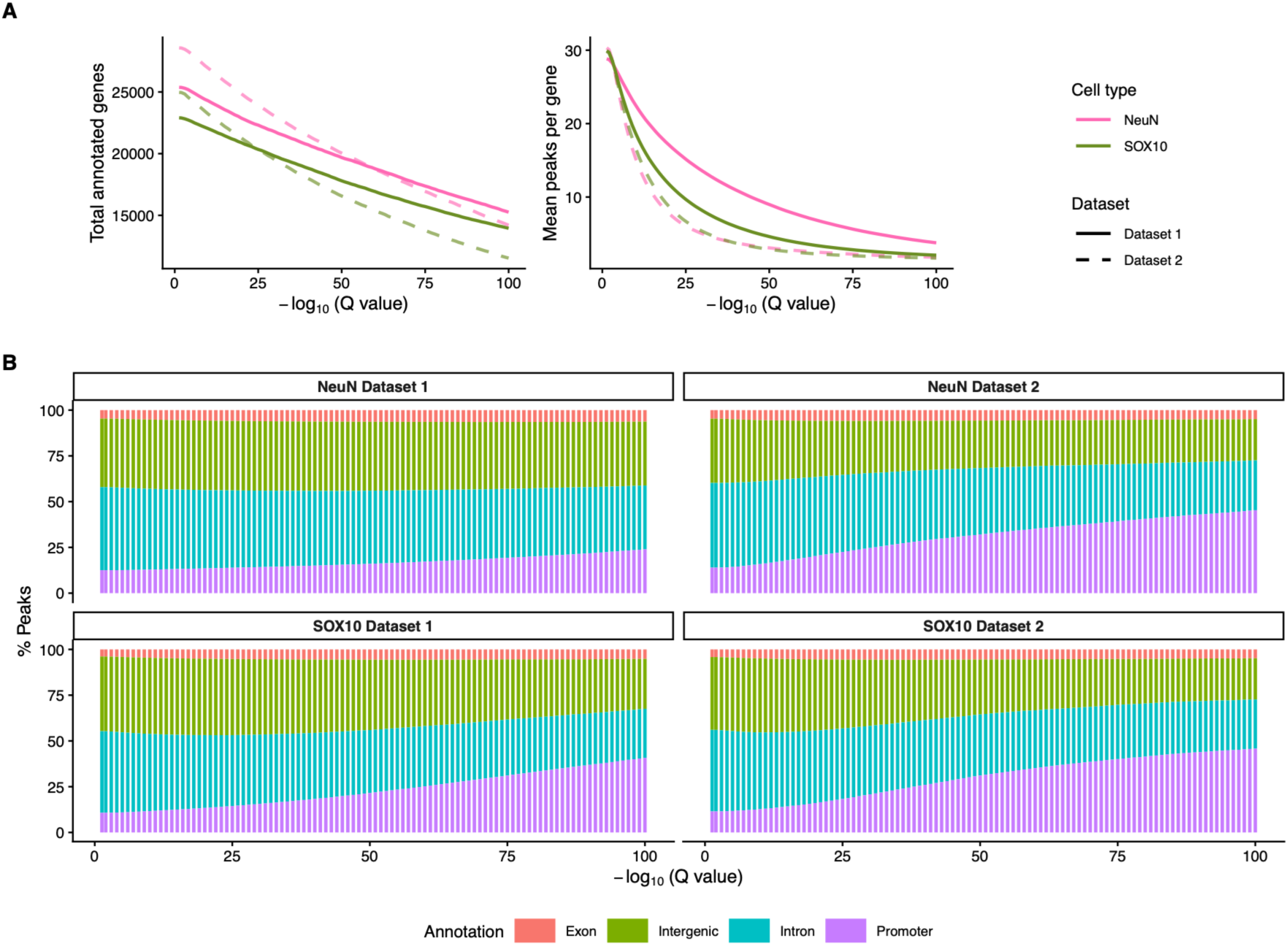
More stringent Q value thresholds change the genomic characteristics of peaks. Annotation statistics for peak sets generated across thresholds ranging from Q<0.05 to 1e-100 are shown for NeuN+ and SOX10+ cell types in both datasets. **A; left panel**) Line graph showing the total number of annotated genes per peak set across thresholds; **A; right panel**) Line graph of mean peaks annotated per gene per peak set across thresholds, where the line colour and style reflect the condition. **B**) Stacked bar charts showing the proportion of peaks assigned to each genomic annotation category dataset and cell type across Q value thresholds.

These observations suggest that the relationship between peak number and Q value comprises two distinct patterns: at more lenient Q values, increasing stringency thresholds removes large numbers of peaks, whereas peak numbers change relatively little at higher thresholds despite further increases in stringency (**Figure 3A**). We hypothesised that the transition between these patterns, i.e. where the gradient of curve becomes noticeably shallower, commonly referred to as the elbow of the curve, reflects the depletion of peaks arising predominantly from background noise and the retention of a more robust set of biologically meaningful peaks. Consequently, we evaluated using segmented regression to identify this breakpoint objectively, providing a data-driven approach that can be applied consistently across diverse datasets to define a suitable Q value threshold (**Figure 5A**).

**Figure 5:**
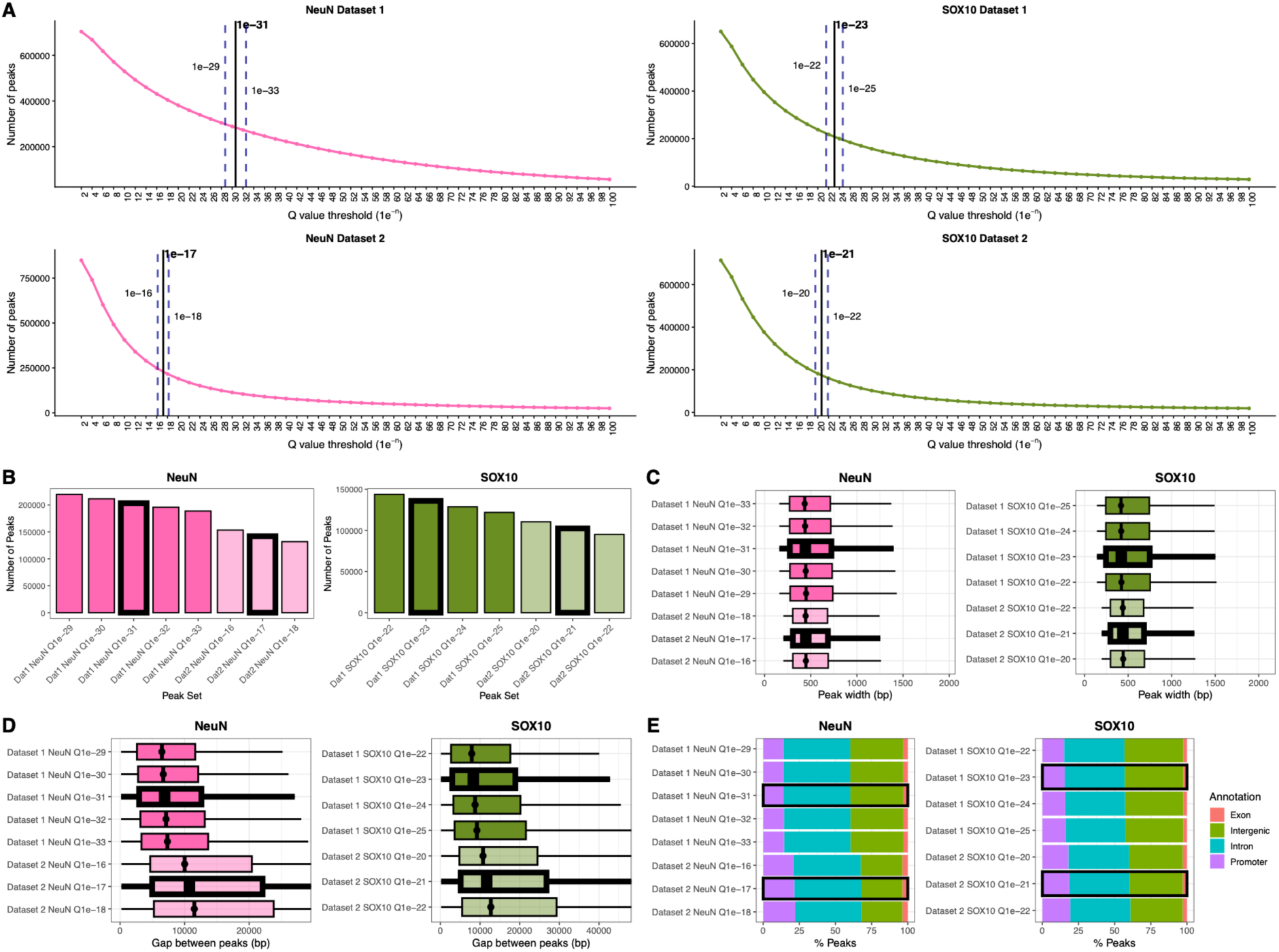
Curated peak sets: more stringent Q value thresholds give more robust peaks sets. **A**) Line graphs showing the number of peaks detected across Q value thresholds. Vertical lines indicate the elbow point identified by segmented regression, which was used to define the curated Q value threshold. Dashed lines show the corresponding 95% confidence intervals (CIs). For panels B-E, all Q value thresholds falling within the 95% CI of the curated threshold were included, whilst the curated threshold is highlighted with black outline. **B**) Bar plots showing the number of peaks detected at Q value thresholds within the 95% CI of the curated threshold. **C**) Boxplots showing the distribution of peak widths across curated peak sets. **D**) Boxplots showing the distribution of interpeak distances across curated peak sets. **E**) Stacked bar plots showing the genomic annotation profiles of curated peak sets.

Applying this approach to both datasets for two cell types (NeuN+ and SOX10+), we found that the optimum threshold tended to be approximately one order of magnitude more significant than the typical default thresholds (Q<0.05), with variability across the cell types and datasets. The most stringent threshold was derived for the NeuN+ fraction in Dataset 1, which was the dataset with the highest sequencing depth. To validate these thresholds, we profiled the characteristics of the resulting “curated” peak sets. Across the four cell type-specific datasets the number of peaks called was of the same order of magnitude (121,835 –195,845 peaks; **Figure 5B**). This represents approximately 3.6-5.5 times fewer than at the default threshold, containing a substantially smaller fraction of the human genome (1.6 × 10^8^-1.8 × 10^8^bp; 5.9-6.7%). In general, these curated peak sets were more likely to be annotated to promoters, and fewer peaks were annotated to each gene. Finally, we compared the peak sets to those calculated with neighbouring significance thresholds within the 95%CI of the elbow to determine how sensitive the results were to the specific threshold. We observed that the properties of peaks in terms of number, width, gaps between neighbouring peaks and annotation proportions were similar across these local alternative thresholds within each dataset and cell types (**Figure 5C-E**). In summary, the characteristics of the curated peak sets were more in line with our expectation, compared to the default significance thresholds.

### More stringent, curated peak sets show higher reproducibility across datasets

A major strength of our study is the availability of two independent cell type-specific datasets generated in the same lab, following the same protocol enabling us to quantify the reproducibility of the cell type-specific peaks. When reproducibility was assessed simply as the proportion of peaks overlapping between datasets, the default peak sets (Q value < 0.05) had a higher overlap (67.42% and 64.21% of Dataset 1 and 2 NeuN+ peaks respectively) compared to the curated peak sets (32.09% and 41.28% of Dataset 1 and 2 NeuN+ peaks, respectively; **Supplementary Figure 13**). However, because the default peak sets covered almost three times more of the genome, they also had a substantially greater probability of overlapping by chance.

Therefore, we used the Forbes coefficient to assess the similarity between two peak sets. This approach quantifies whether they overlap more than expected by chance, given how much of the genome they cover.^50^ Comparing pairwise the four conditions, we found that the curated peak sets showed 7.90-18.1 times the overlap expected by chance, while default peak sets showed only 1.80-2.66. ^50^ The strongest degree of overlap was observed between the same cell fraction across different datasets, aligning with the cell-specificity of the data (**Figure 6A**). The overlap was strengthened when limiting the comparison to peaks annotated to promoters (74.2 and 127 for NeuN+ and SOX10+ peaks respectively) (**Figure 6A**). In addition, we used the per-peak-pair Jaccard metric to quantify the extent of overlap between two peaks while accounting for their respective length. Despite a lower proportion of peaks that overlap between datasets in the curated peak sets, the Jaccard scores were higher on average for both cell types, with mean overlap of 56% in curated NEUN+ peak set compared to 38% for default NeuN+ peak set and 56% compared to 44% in SOX10+ curated and default peak sets respectively (**Figure 6B**). For both cell types, filtering the peaks to those detected in both datasets dramatically increased the proportion annotated to the promoters compared to those specific to one dataset. With the strongest enrichment when intersecting the curated peak sets (**Figure 6C**). Furthermore, a higher percentage of genes annotated to peaks which were detected in the curated peak sets of both datasets were classed as brain expressed (**Supplementary Figure 14**). Finally, we extended the application of our elbow method to determine a robust significance threshold for calling peaks in the other cell fractions, with fewer samples (IRF8+ and TN). As expected, there was a clear linear relationship between the curated peak set Q value and sequencing depth (p<1e^-3^), further validating the approach in both large and small datasets (**Figure 7**).

**Figure 6:**
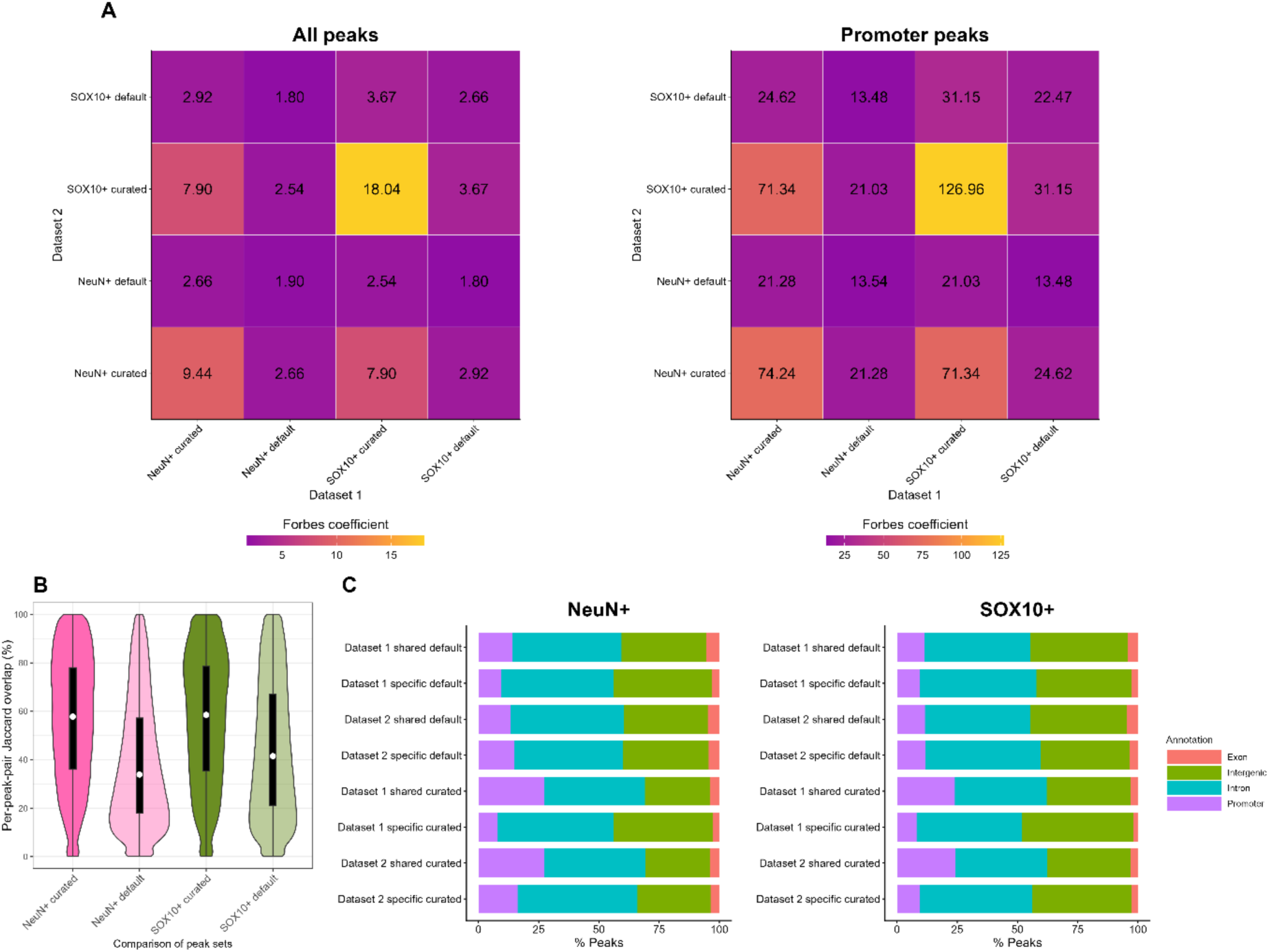
Curated peak sets are more reproducible across datasets. **A**) Heatmap showing Forbes coefficients quantifying the overlap between peak sets across datasets and cell types for the default peak sets (Q < 0.05) and curated peak sets (Q < threshold determined by elbow analysis). The left heatmap includes all peaks, whereas the right heatmap is restricted to promotor-annotated peaks. **B**) Violin plots showing the distribution of pairwise peak-level Jaccard overlap between datasets for each cell type, comparing the default and curated peak sets. **C**) Stacked bar plots showing the percentage of peaks assigned to each genomic annotation categories for peak shared or specific to each dataset for the default peak sets (Q < 0.05) and curated peak sets (Q < threshold determined by elbow analysis).

**Figure 7:**
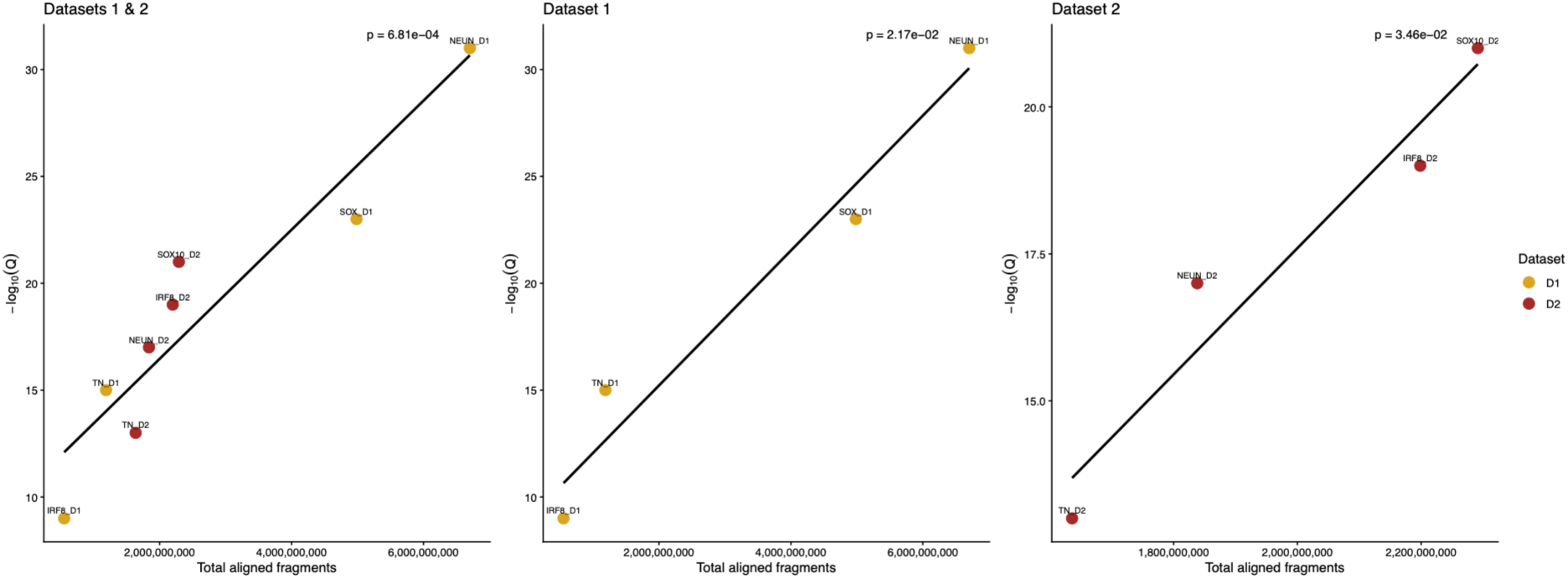
Relationship between sequencing depth and curated peak calling thresholds across Dataset 1 and Dataset 2, for all four cell fractions. Scatter plots show the relationship between total aligned fragments and curated peak calling significance thresholds, expressed as -log₁₀(Q), for Dataset 1 (D1, yellow) and Dataset 2 (D2, brown). The left panel shows both datasets combined, while the middle and right panels show Dataset 1 and Dataset 2, respectively. Each point represents a cell-type-specific peak calling threshold used to generate the curated peak sets. Black lines indicate linear regression fits for the association of sequencing depth with -log₁₀(Q value), with the corresponding p values for the regression coefficients reported in each panel.

## Discussion

We present a novel pipeline for integrative peak calling in population scale ATAC-seq studies: PIPPA, validated across two independent cell-sorted datasets, with a data-driven strategy to address inflation in peak calling observed in deeply sequenced datasets. This work provides the foundations for future epidemiological studies of open chromatin both in large studies of bulk tissue and nuclei-sorted samples.

Although ATAC-seq is now widely used, many bioinformatic workflows still rely on software and tools originally developed for other epigenomic assays. The field is shifting from bulk-tissue analyses to cell type-specific approaches, including profiling sorted populations and single-nuclei ATAC-seq, to improve the biological interpretation of chromatin accessibility differences. We developed PIPPA to reproducibly automate the quality control of large-scale nuclei-sorted or bulk chromatin accessibility datasets. Building on our established framework for preprocessing cell-specific DNA methylation data^12,13^, the pipeline takes raw sequencing reads through complementary stages assessing sequencing and library quality, sample identity, and cell type identity (**Supplementary Figure 1**). Consistent with other epigenomic data, we found that the major axes of variation strongly reflects cell type identity, a feature that we exploit in our pipeline. As with DNA methylation data, this potentially confers benefits for statistical power in epidemiology studies, meaning cell sorting can be strategic where samples are limited.

Defining a consensus peak set is critical for downstream analyses, requiring a balance between maximising biological signal and minimizing artefactual peaks that obscure true phenotypic associations. We generated a consensus peak set by calling peaks on a combined signal created from all aligned reads merged into a single bam file. Peak calling algorithms have historically been developed and evaluated using relatively modest numbers of samples, where true enrichment is expected to be clearly distinguishable from background signal. In contrast, our analyses showed that standard software performs particularly poorly on very deep datasets, with the number of detected peaks continuing to rise at default Q value thresholds, rather than converging to an upper bound. This phenomenon, often referred to as “peak saturation” has been reported for other sequencing assays^51^, and is evident in existing ATAC-seq studies that report unexpectedly large numbers of peaks. As sequencing depth increases, read coverage accumulates in regions flanking the core signal, extending peak boundaries and resulting in additional or broader regions being called as peaks. This not only leads to an unexpected number of peaks, it also influences the characteristics of peaks called, with an abundance of narrow peaks close together annotated to virtually every gene and with increasing number of peaks per gene. At default thresholds there is limited replication of peaks across independent datasets, fitting with the idea of spurious peaks across the genome erroneously included in the peak set. These results suggest that the largest peak set is not necessarily the most biologically informative, particularly in deeply sequenced datasets.

To mitigate the risk of false positive peaks underpinning downstream analyses, we propose using a more stringent, data-driven Q value threshold. Specifically, our approach uses the elbow of the peak number versus Q value curve to identify a dataset-specific threshold. This method provides a confidence interval which provides a range of values to choose from, depending upon the objectives of the study. A conceptually similar cut-off analysis is available in MACS, in which multiple thresholds are tested and the resulting number of peaks, as well as their average and total lengths, are reported. However, the lowest Q value threshold tested by MACS is 1e^-10^, which, as we observed, is unlikely to be sufficiently stringent for datasets with higher sequencing depth. Using two large datasets we show that across well-powered studies, we observe substantially different numbers of peaks at equivalent thresholds. In addition, we show heterogeneity in the rate at which peaks are called, highlighting the variation in efficacy of ATAC-seq datasets likely due to differences in experimental quality. Using our approach to determine a dataset specific threshold, numerically comparable peak sets per cell fraction can be called across datasets of different statistical properties.

These findings also complicate the guidance on appropriate sequencing depth targets. Sequencing depth must provide sufficient power for reliable peak detection while also providing sufficiently sensitive quantification to facilitate downstream analyses such as differential accessibility testing. In our pipeline, we adopt the ENCODE recommendation of at least 25 million unique aligned read pairs (50 million reads) per sample, as a practical threshold in the absence of any empirical data. Importantly, the sequencing depth required to comprehensively define a regulatory landscape may differ from that needed to identify robust quantitative differences between samples.

Whilst we show that this curated threshold is better than the standard thresholds, it may be that there are other ways to derive this threshold depending on the goals of the study and what the peak set will be used for. As an alternative we could model the Q value distribution as a mixture of two distributions, such that we can then subtract the inflated distribution. This requires the full Q value distribution to be calculated, a computationally expensive and not straightforward ambition for ATAC-seq data given the correlation between neighbouring sites. It requires subdividing the genome into bins, but determining the size of these bins is non-trivial. Such an approach has been proposed for ChIP-seq data but relies on both IP enriched and control samples to be sequenced.^52^

There is also substantial variation in how consensus peak sets are generated across multi-sample ATAC-seq studies. A common alternative approach is to find consensus peaks, by merging and taking the union across those called at individual sample level. This approach does not utilise the power of the full dataset but given that each individual sample will have lower number of reads, arguably reduces the inflated Q value limitation described here. Another emerging approach involves taking the peaks called at a standard threshold from the combined (merged) samples as a reference, then sub-setting these according to peaks found in *n* individual samples. However, this approach is largely dependent on the sensitivity of the sample with the maximum coverage and the appropriateness of the Q value applied for the sequencing depth of the individual sample(s). It is arguable that peak calling could in time become redundant as once there are sufficient datasets, the total set of peaks can be defined and all subsequent datasets evaluated against these. While this may prevent the identification of artefacts in noisy datasets, it rests on the assumption that a stable peak set exists overlooking context-specific peaks arising from differences in cell state, response to stimuli, ageing or other exposures.

A major strength of our study is the availability of two large, cell sorted ATAC-seq datasets, generated within the same laboratory following the same experimental protocol. This enabled us to evaluate the reproducibility of our peak sets, providing an independent assessment of the proposed methodology. However, we recognise that access to a second dataset is a unique scenario; our pipeline was therefore designed to function without one. Given the propensity for inflated peak sets in large-scale ATAC-seq datasets, we recommend that peak sets are validated where possible. More broadly, our findings reinforce the importance of validating epidemiological associations, with validating the true existence of a peak a critical first step, prior to confirming differences detected at those peaks.

Our study should be considered in light of the following limitations. Given the widespread use of MACS for peak calling, we limited our comparisons to this software and the typical parameter choices and did not consider alternative peak calling tools developed specifically for ATAC-seq such as Genrich^53^, HMMRATAC^33^ and Replicative contrastive learner (RCL)^55^. A comprehensive comparison of these methods in large, deeply sequenced datasets would be valuable, particularly because most previous benchmarking has been undertaken in substantially smaller datasets. It is unclear whether these methods also are afflicted by inflated peak sets, and it may be that our approach to derive an appropriate threshold could transfer to other peak callers. In addition, we have limited our study to FANS-isolated nuclei derived from post-mortem brain tissue. Given the challenges with getting high-quality DNA from these samples, our data may be affected by higher technical noise than bulk tissue. Consequently, our findings may be exaggerated relative to studies of bulk tissue.

In summary, we introduce PIPPA an automated and scaleable pipeline for processing and QC of ATAC-seq data. We demonstrate that dataset-specific, stringent significance thresholds are needed to generate reproducible peak sets. We propose a novel approach for curating peaks that facilitates more reliable downstream analysis, establishing a framework for future epidemiological studies focused on identifying epigenetic variation associated with development and disease.

## Acknowledgements

We acknowledge the supply of samples from Brains for Dementia Research and Autism BrainNet. For the purpose of open access, the author has applied a CC BY public copyright license to any Author Accepted Manuscript version arising from this submission.

## Author contributions

EH and JM conceived and supervised the study. ED supervised the lab work, BC and JB performed the lab work; EH designed and coded the initial version ATAC-seq BrainFANS pipeline, MFP and PW refined the pipeline and MFP curated the publicly available scripts with supervision from EH. EH, PW, MFP, JM and AB collaborated for novel methods development. MFP and PW analysed and visualised the data. PW, MFP, EH & JM wrote the manuscript, with contributions from all authors.

## Consortia

The members of the Youth-GEMs Consortium are Bart Rutten, Sinan Gülöksüz, Therese van Amelsvoort, Lotta-Katrin Pries, Bochao Danae Lin, Angelo Arias-Magnasco, Erika van Hell, Mary Cannon, David Cotter, Melanie Föcking, Subash Raj Susai, Elisabeth Binder, Jim van Os, Jeroen Pasterkamp, Anna Wiersema, Marco Boks, Winni Schalkwijk, Karim Lekadir, Esmeralda Ruiz Pujadas, Noussair Lazrak, Ian Kelleher, Jenni Leppänen, Valentina Kieseppä, Simona Karbouniaris, Lisette van der Poel, Mariël Kanne, Marijke Kolk, Hanske Douwenga, Jordi Rambla, Arcadi Navarro, Liina Nagirnaja, Aldar Cabrelles Munoz, Lauren A. Fromont, Jaanus Harro, Triin Kurrikoff, Reigo Reppo, Dejan Stevanovic, Aleksa Milevic, Jasna Jancic, Marija Nikolic, Maria Bulgheroni, Laura Giani, Margherita La Gamba, Tomislav Franic, Mia Plenkovic, Covadonga M. Diaz-Caneja, Celso Arango, Marta Ferrer-Quintero, Renzo Abregú-Crespo, Emily Guerra-Blacio, Nuria Martín-Martínez, Christel Middeldorp, Enda M. Byrne, Swathi Hassan Gangaraju, Sushma Marla, Jonathan Mill, Eilis Hannon, Emma Dempster, Philippa M. Wells, Robin Murray, Andrea Danese, Alexander L. Richards, Lucy Riglin, and Michael C. O’Donovan.

## Data availability

Code for the analysis and visualisations presented in this manuscript are available at https://github.com/EpigeneticsExeter/PIPPAmanuscript

The ATAC-seq pipeline (PIPPA) can be found at: https://github.com/ejh243/BrainFANS/tree/master/sequencing/ATACSeq

