## Supplementary Material for "Best Practices for Pre-Processing Large-Scale ATAC-Seq Data: Why Epidemiological Studies Demand Greater Stringency"

### PIPPA: Supplementary Material

### Tables

#### Supplementary Table 1: Summary of sample demographics and quality control

| | Dataset 1 | Dataset 2 |
|---|---|---|
| N (NeuN+; SOX10+; IRF8+; TN) | 109;91;5;17 | 34;31;25;30 |
| Age (*range*) | 60-90 | 6-91 |
| QC1: Fail Alignment Rate (<80%) n(%) | 15 (3%) | 0 (0%) |
| QC1: Fail Aligned Reads (<50million) n(%) | 72 (16%) | 31 (18%) |
| QC1: Fail NRF(<0.7); n(%) | 11 (2%) | 18 (10%) |
| QC1: Fail PBC 1 (<0.7;) n(%) | 11 (2%) | 18 (10%) |
| QC1: Fail PBC2 (<1); n(%) | 0 (0%) | 0 (0%) |
| QC1: Fail Nuc-Free (<0.25); n(%) | 0 (0%) | 0 (0%) |
| QC1: Fail Mono-Nuc (<0.2); n(%) | 134 (30%) | 3 (2%) |
| QC2: Fail Genotype Check; n(%) | 66( 15%) | NA* |
| QC2: Fail Sex Check; n(%) | 16 (4%) | 0 (0%) |
| QC3: Fail cell type check; n(%) | 13 (6%) | 2 (4%) |

**Genotype data unavailable to facilitate the genotype check in Dataset 2*

### Supplementary Figures

#### Supplementary Figure 1

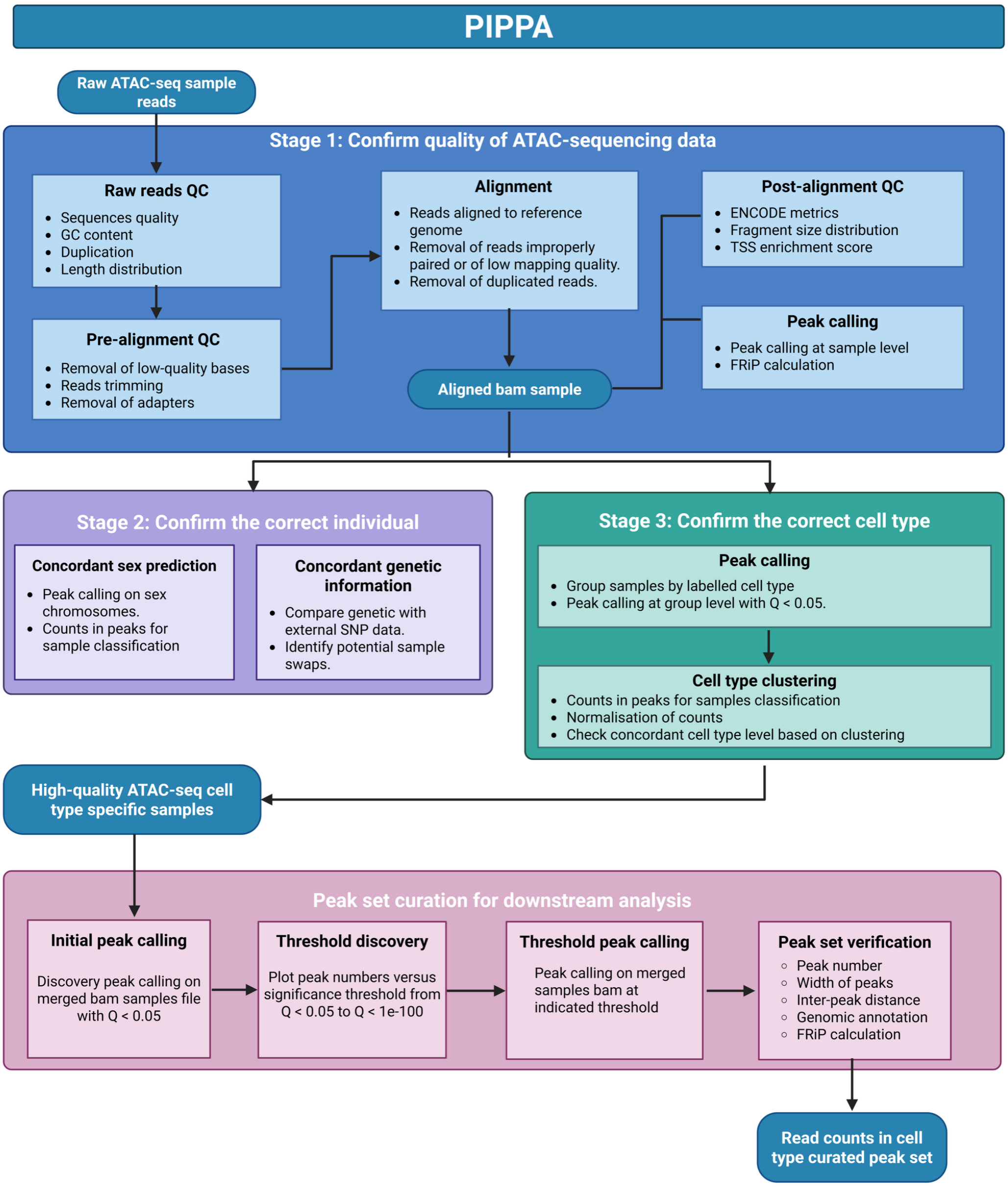

**Supplementary Figure 1: PIPPA overview.** The pipeline consists of three sequential quality-control stages to confirm: 1) ATAC-sequencing data quality, 2) sample identity and 3) cell type label. These checks are followed by the generation of a curated peak set for downstream analyses.

### Supplementary Figure 2

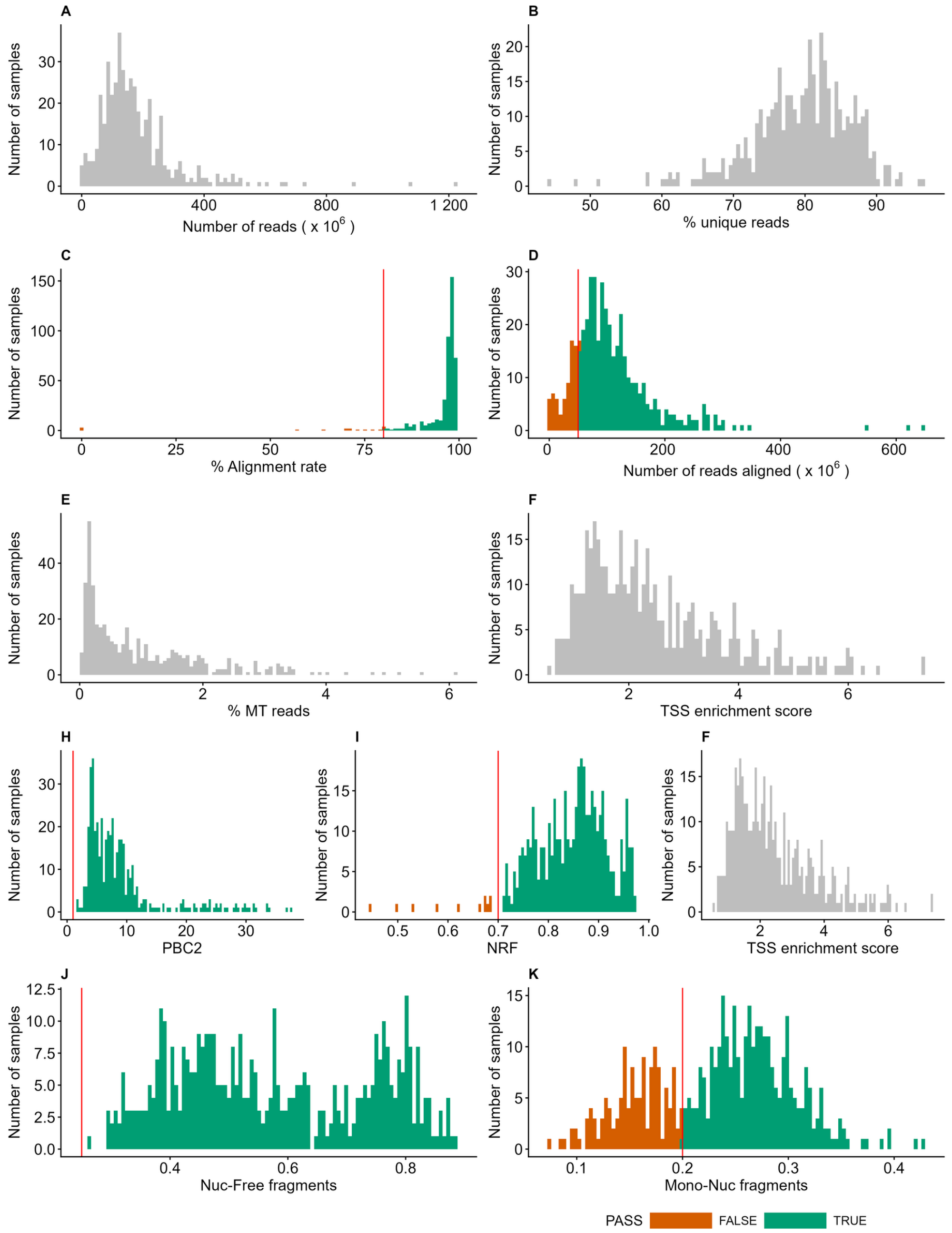

**Supplementary Figure 2: PIPPA stage 1 summary.** Distributions of data properties and QC metrics used to identify high-quality ATAC-seq datasets as part of Stage 1 of the BrainFANS pipeline. Shown are histograms of: **A**) number of reads before QC; **B**) percentage of unique paired-end reads before QC; **C**) alignment rate; **D**) number of aligned reads after post-alignment filtering; **E**) percentage of mitochondrial reads; **F**) TSS enrichment score; **G–I**) ENCODE QC metrics including PCR bottlenecking coefficients PBC1 (**G**) and PBC2 (**H**), and the non-redundant fraction (NRF, **I**); **J–K**) fragment size distribution metrics showing the proportion of nucleosome-free regions (Nuc-Free, **J**) and mono-nucleosome fragments (Mono-Nuc, **K**). Data shown are for Dataset 1. Where QC metrics were used to filter samples, applied thresholds are indicated by vertical red lines. Histogram colours denote whether samples at each value passed or failed the corresponding QC metric.

### Supplementary Figure 3

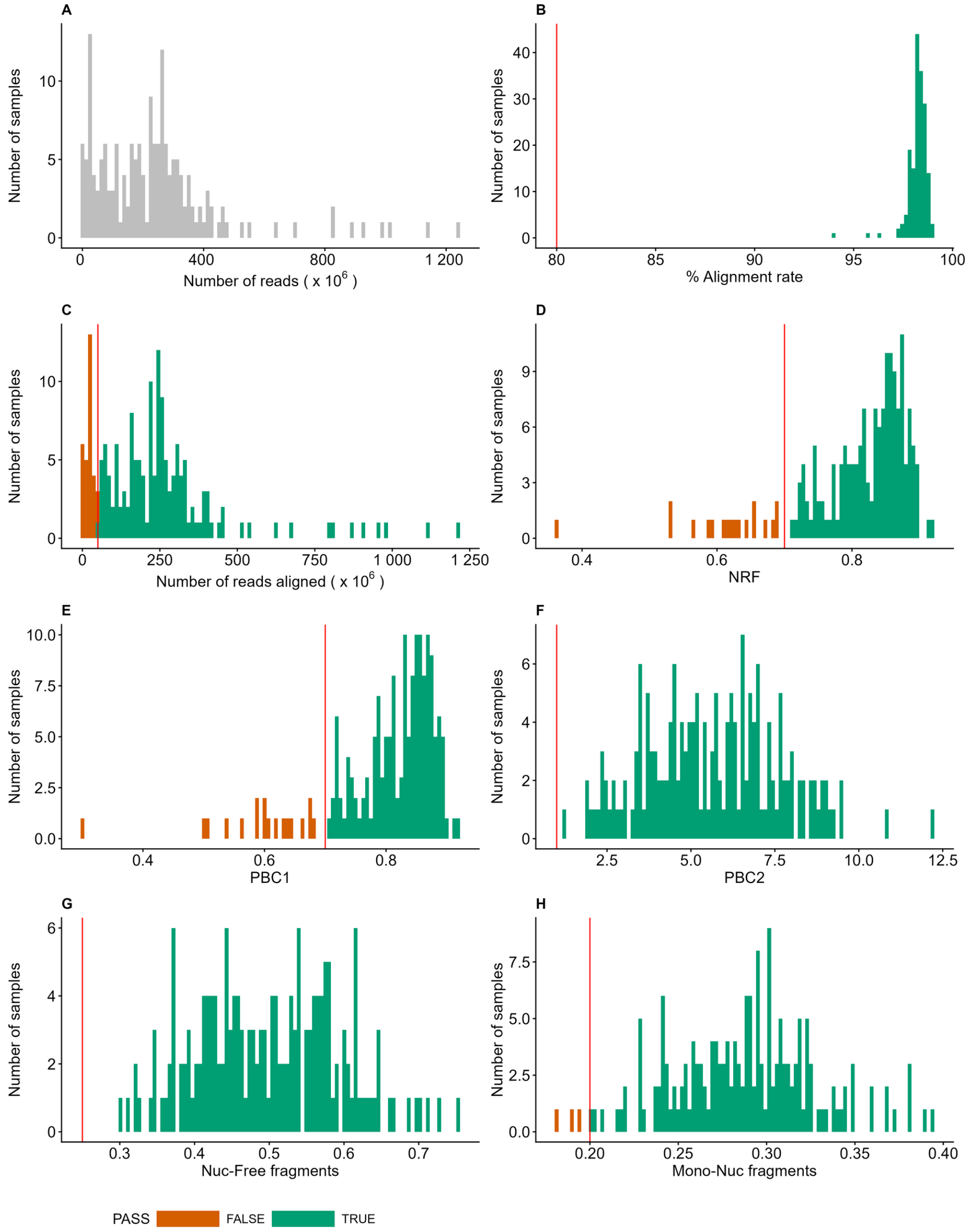

**Supplementary Figure 3: PIPPA stage 1 summary.** Distributions of data properties and QC metrics used to identify high-quality ATAC-seq datasets as part of Stage 1 of the BrainFANS pipeline. Shown are histograms of: **A**) number of reads before QC; **B**) alignment rate; **C**) number of aligned reads after post-alignment filtering; **D–F**) ENCODE QC metrics including PCR bottlenecking coefficients PBC1 (**D**) and PBC2 (**E**), and the non-redundant fraction (NRF, **F**); **G–H**) fragment size distribution metrics showing the proportion of nucleosome-free regions (Nuc-Free, **G**) and mono-nucleosome fragments (Mono-Nuc, **H**). Data shown are for Dataset 2. Where QC metrics were used to filter samples, applied thresholds are indicated by vertical red lines. Histogram colours denote whether samples at each value passed or failed the corresponding QC metric.

### Supplementary Figure 4

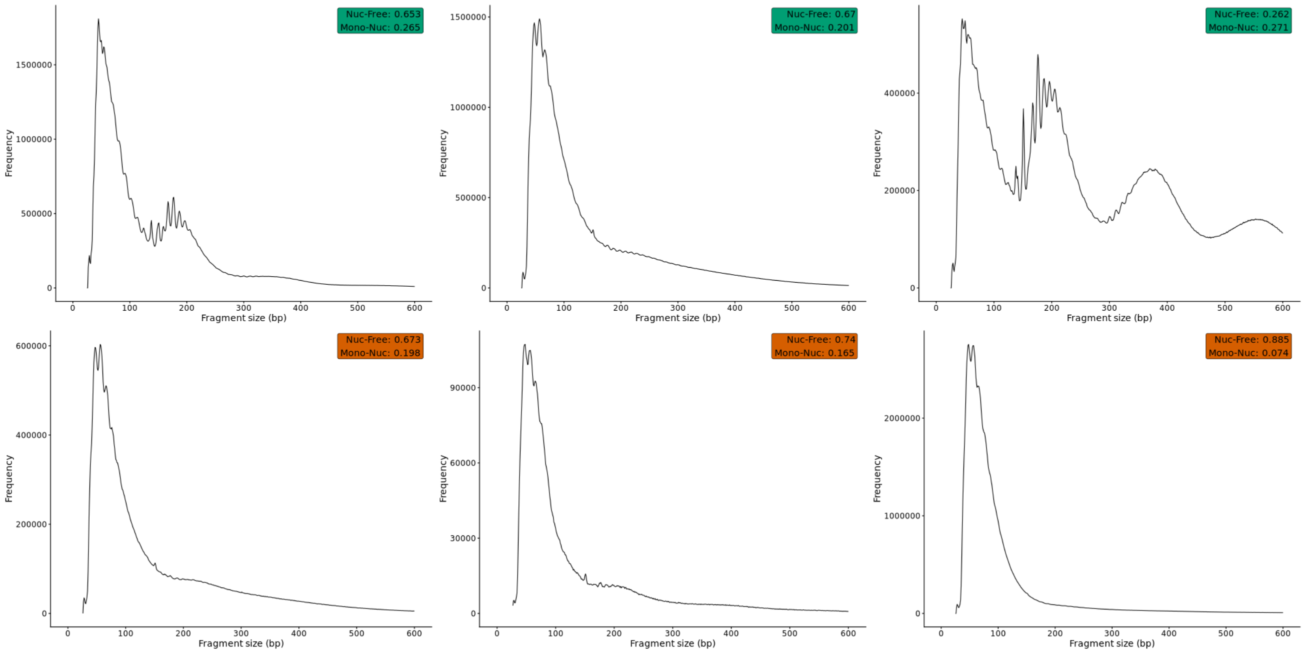

**Supplementary Figure 4: Fragment size distribution.** Examples of fragment size distribution of samples that pass (upper row) or fail (bottom row) the proportion of nucleosome-free regions (Nuc-Free) or mono-nucleosome fragments (Mono-Nuc). Colours indicate pass or fail.

#### Supplementary Figure 5

**A**

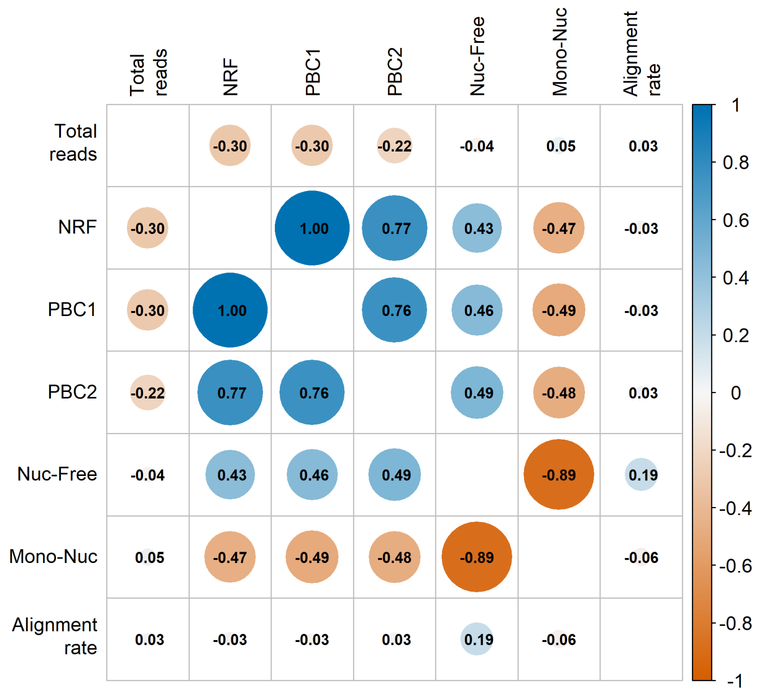

**B**

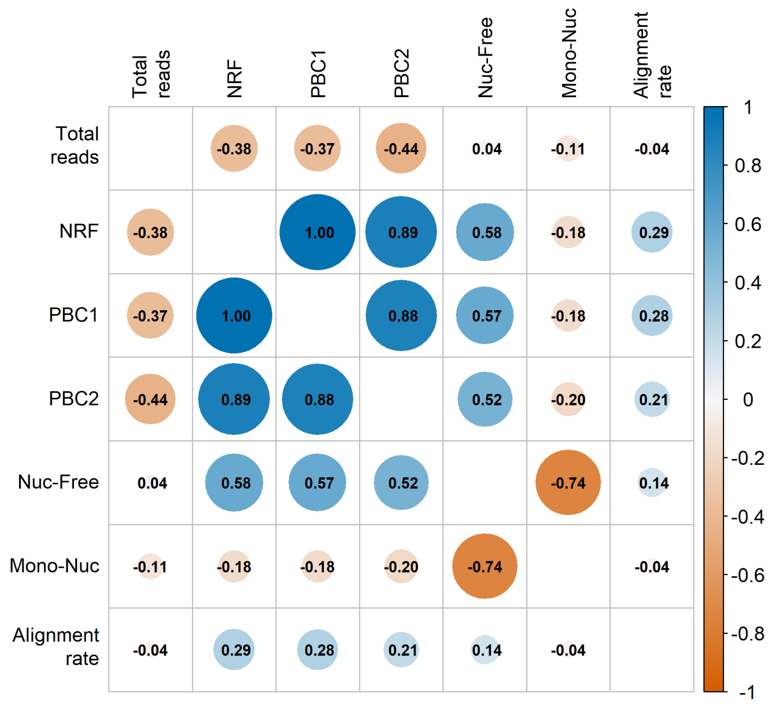

**Supplementary Figure 5: Correlation of QC metrics used in Stage 1 of the PIPPA to confirm successful ATAC-seq profiling.** Matrix of Pearson correlation statistics between all pairs of QC metrics. Circle size reflects the magnitude of the correlation coefficient, and circle colour indicates direction and strength, with representing positive correlations and red representing negative correlations. **A)** Data shown for Dataset 1. **B**) Data shown for Dataset 2.

#### Supplementary Figure 6

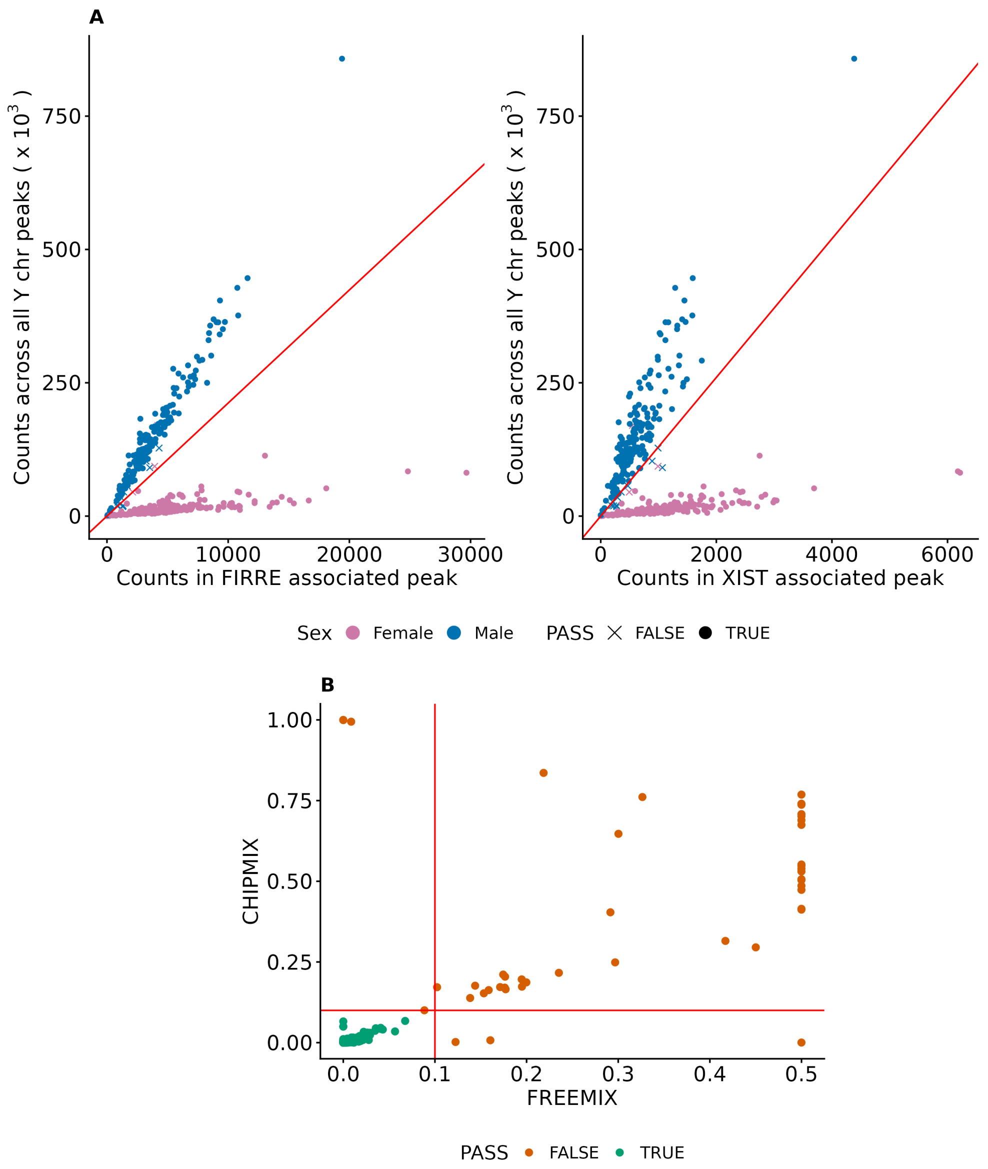

**Supplementary Figure 6: Stage 2 of the PIPPA to confirm sample donor identity. A**) Scatterplot showing sex-verification metrics, comparing read counts in the nearest upstream peak to FIRRE (left) and XIST (right) genes (x-axis) against total read counts in peaks across the Y chromosome (y-axis). Point colour indicates the labelled sex of each sample, and point shape denotes whether the predicted sex matches the labelled sex. Red line indicates threshold to predict sex where samples above the line are predicted as male and those below as female. **B**) Scatterplot of genotype verification metrics, where FREEMIX (x-axis) reflects the probability of contamination and CHIPMIX (y-axis) reflects the probability that a sample matches a different donor.  Each point represents a sample, coloured according to whether it fails either metric. Applied QC thresholds are shown as red lines. Data shown are for Dataset 1. Data shown for Dataset 1.

#### Supplementary Figure 7

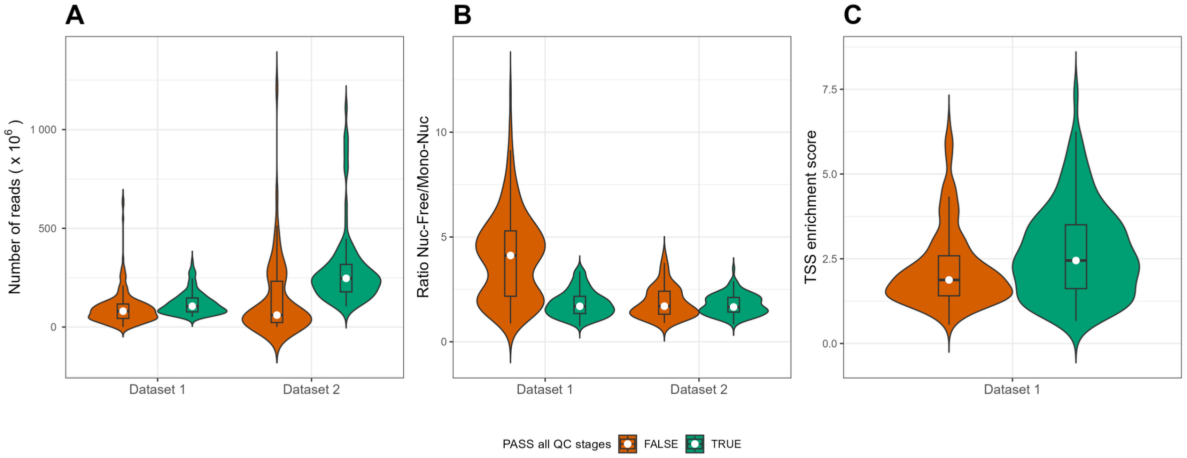

**Supplementary Figure 7:** Violin plots showing the distribution of **A**) sequencing depth (number of reads; y-axis), **B**) nucleosome-free to mono-nucleosome ratio (Nuc-Free/Mono-Nuc; y-axis) and **C**) TSS enrichment score (y-axis) for samples that pass or fail all QC stages. Colours indicate whether sample has passed all QC stages.

#### Supplementary Figure 8

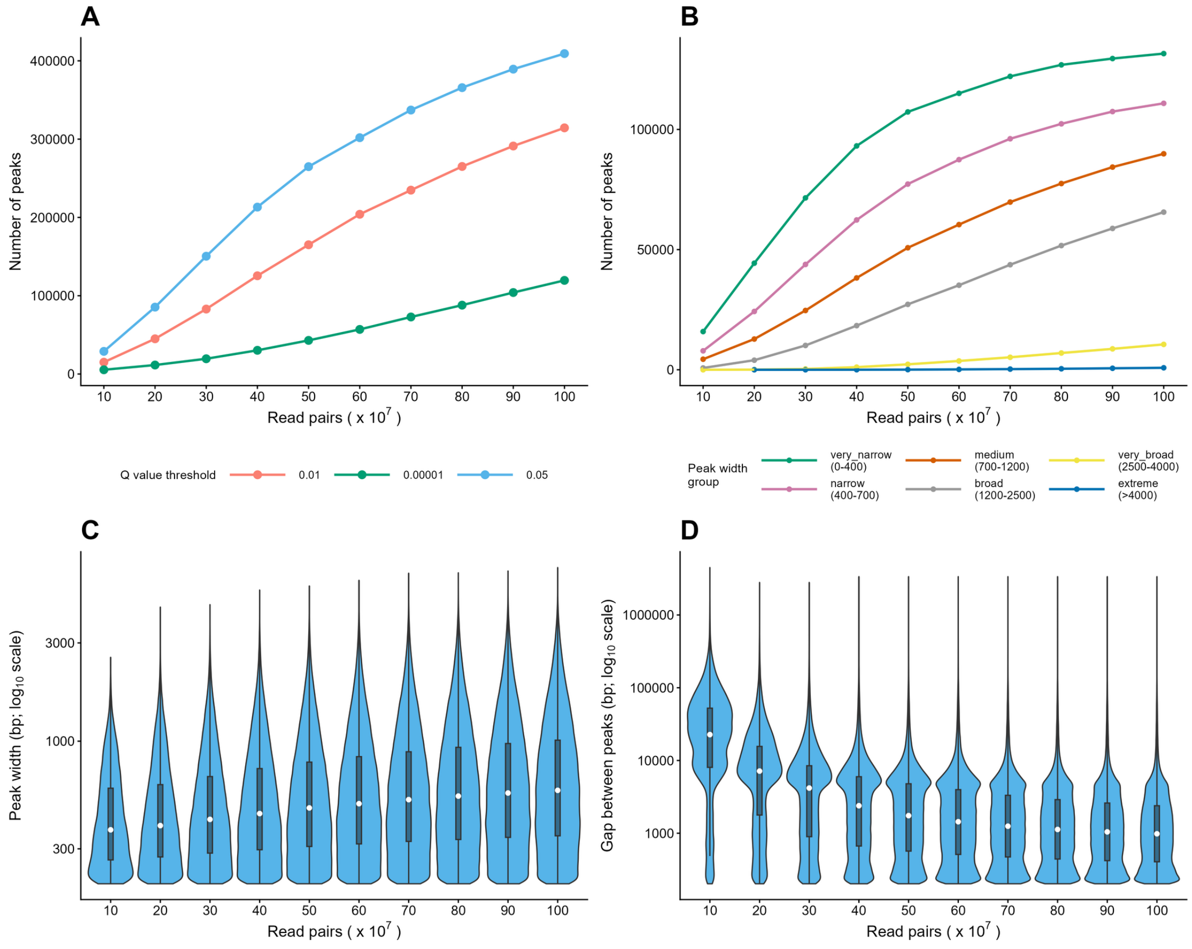

**Supplementary Figure 8: peak calling exhibits sequencing-depth-dependent changes in peak characteristics. A**) Line graphs showing the number of peaks (y-axis) across increasing sequencing depths (number of read pairs; x-axis), with each line represents a different Q value threshold. **B**) Line graphs showing the number of peaks (y-axis) identified at increasing sequencing depths (x-axis), where each line represents the peaks grouped by width. Violin plots showing the distribution of **C**) peak width (bp) and **D**) interpeak distance (bp) for peaks called at increasing amounts of sequencing depth. Peaks were called in Dataset 1 SOX10+ samples only; for B-D peaks were called at a Q < 0.05 threshold.

#### Supplementary Figure 9

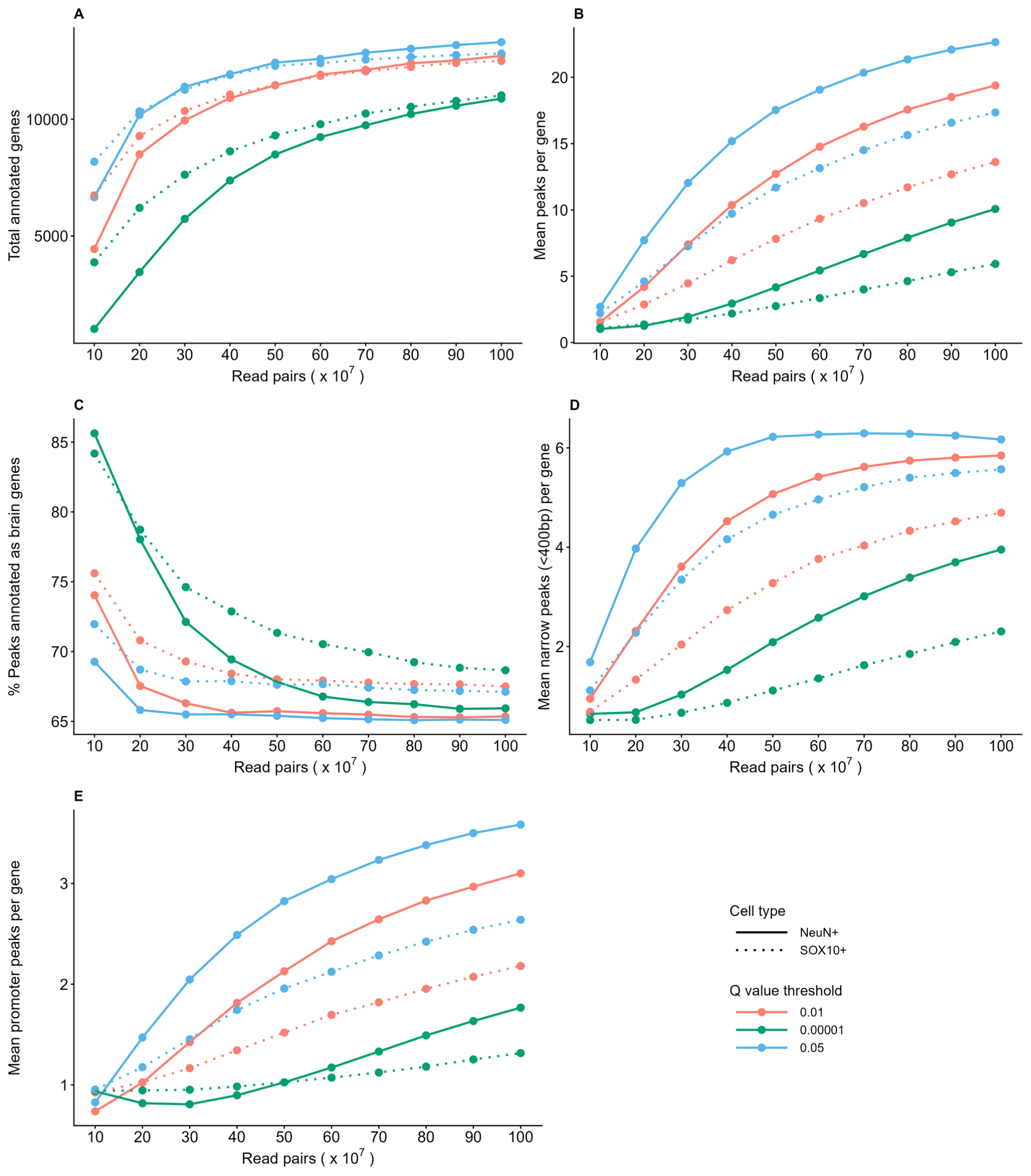

**Supplementary Figure 9: Genomic distribution of "inflated" peak sets at default thresholds.** Line graphs showing sequencing depth (number of read pairs; x-axis) against **A**) the number of annotated genes (y-axis); **B**) the mean number of peaks annotated per gene (y-axis); **C)** the proportion of annotated genes classified as brain genes (y-axis); **D**) the mean number of narrow peaks (< 400bp) annotated per gene (y-axis) and **E**) the mean number of promoter-annotated peaks per gene. Line colours indicated the Q value threshold used for peak calling and line style denotes the cell type. All peaks were called in Dataset 1 NeuN+ and SOX10+ samples respectively. Only protein-coding genes have been considered here.

#### Supplementary Figure 10

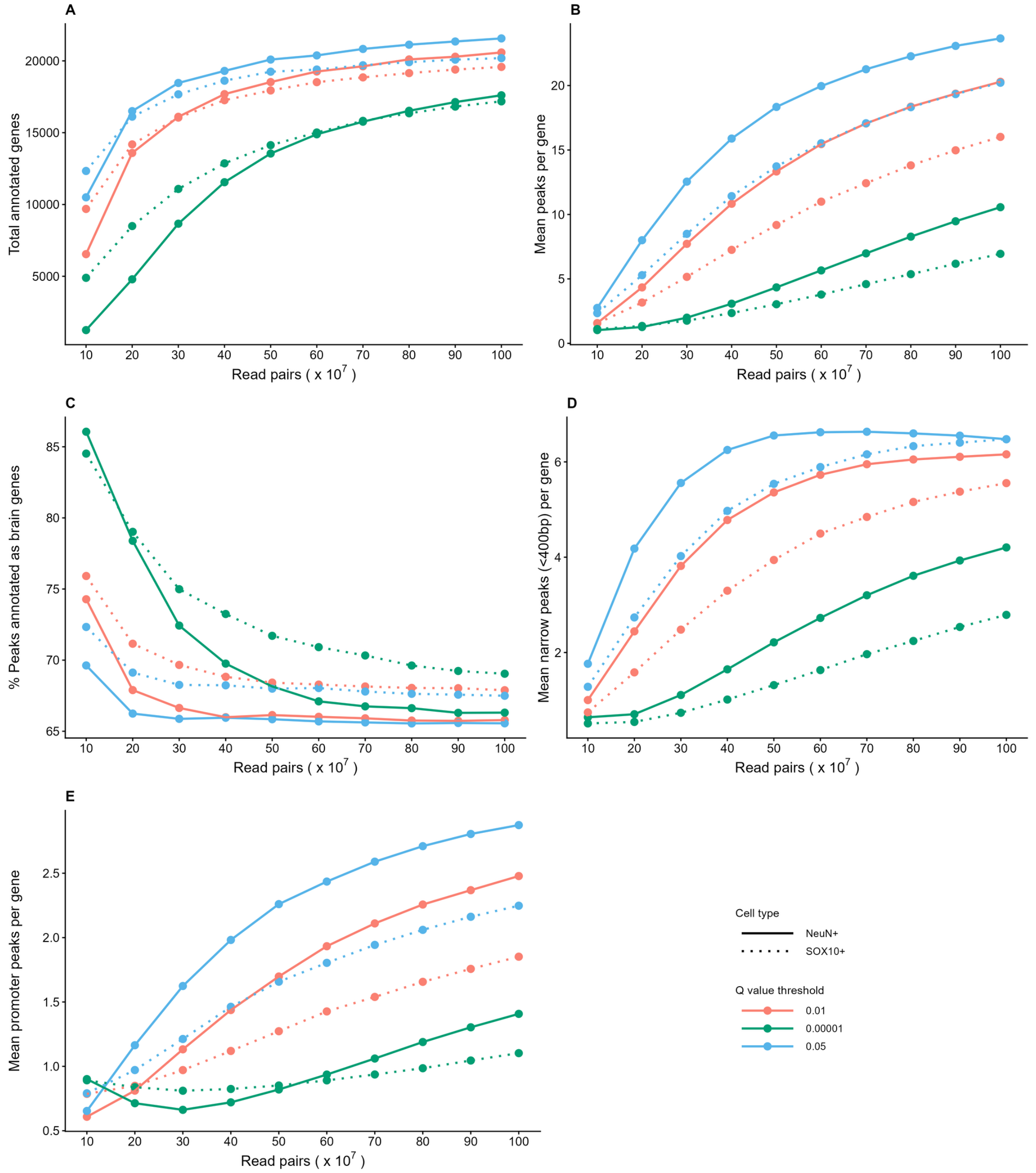

**Supplementary Figure 10: Genomic distribution of "inflated" peak sets at default thresholds.** Line graphs showing sequencing depth (number of read pairs; x-axis) against **A)** the number of annotated genes (y-axis); **B**) the mean number of peaks annotated per gene (y-axis); **C)** the proportion of annotated genes classified as brain genes (y-axis); **D**) the mean number of narrow peaks (< 400bp) annotated per gene (y-axis) and **E**) the mean number of promoter-annotated peaks per gene. Line colours indicated the Q value threshold used for peak calling and line style denotes the cell type. All peaks were called in Dataset 1 NeuN+ and SOX10+ samples respectively.

#### Supplementary Figure 11

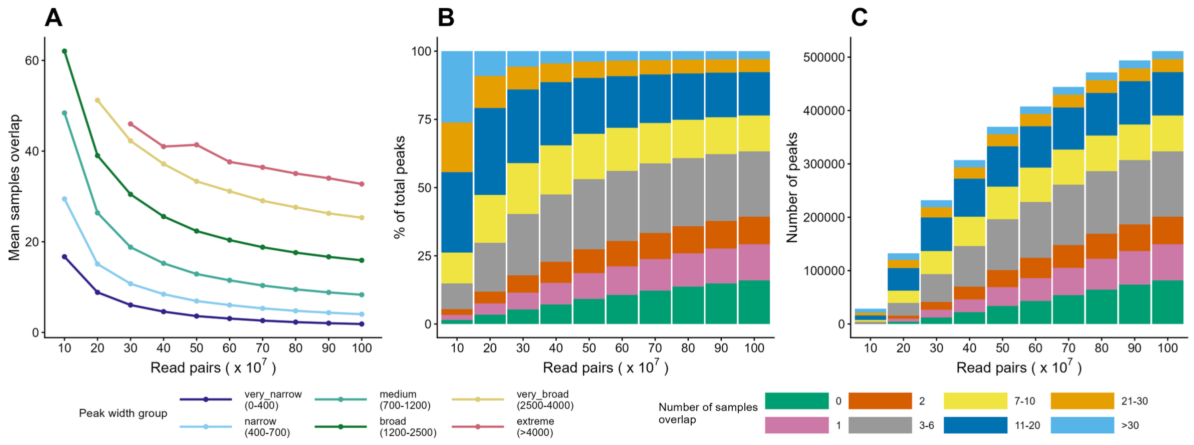

**Supplementary Figure 11: Filtering peak sets based on sample-level peaks is biased. A**) Line graph of the mean number samples each sample overlaps (y-axis) for peak sets at increasing sequencing depth (x-axis), where lines represent the peaks width. Stacked bar plots of the **B**) percentage of peaks (y-axis) and **C**) number of peaks (y-axis) at increasing sequencing depth (x-axis), where peaks are grouped by the number of samples they overlap with. Panels show data for Dataset 1 cell type NeuN+ peak sets called with Q value 0.05.

### Supplementary Figure 12

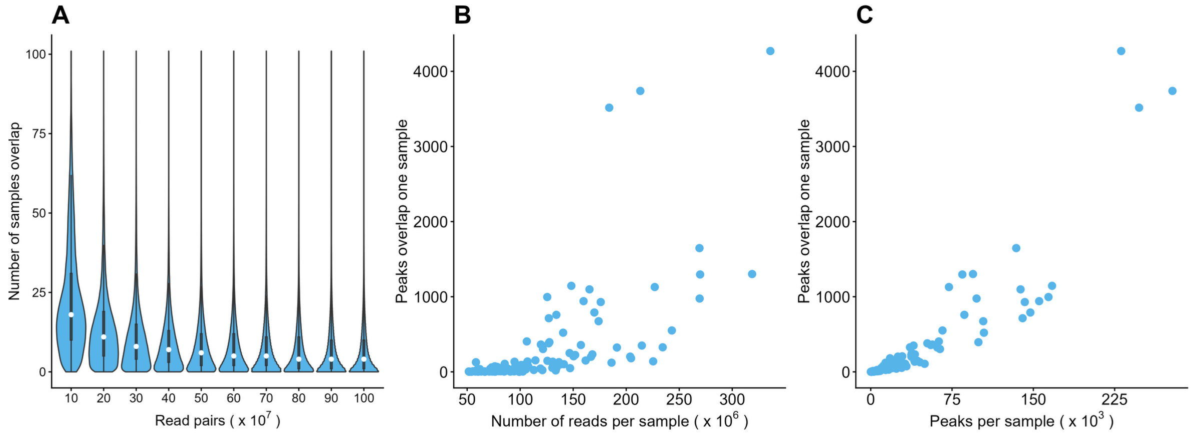

**Supplementary Figure 12: The overlap between merged and sample-level peaks is influenced by sample-level data quantity. A**) Violin plots showing the distribution of the number of samples overlapping each peak (y-axis) for peak sets generated at increasing sequencing depths (x-axis). **B**) Scatterplots showing the relationship between the number of peaks in the merged peak set that overlap each individual sample and the number of read pairs in that sample. **C**) Scatterplots showing the relationship between the number of peaks in the merged peak set that overlap each individual sample and the total number of peaks detected in that sample.

### Supplementary Figure 13

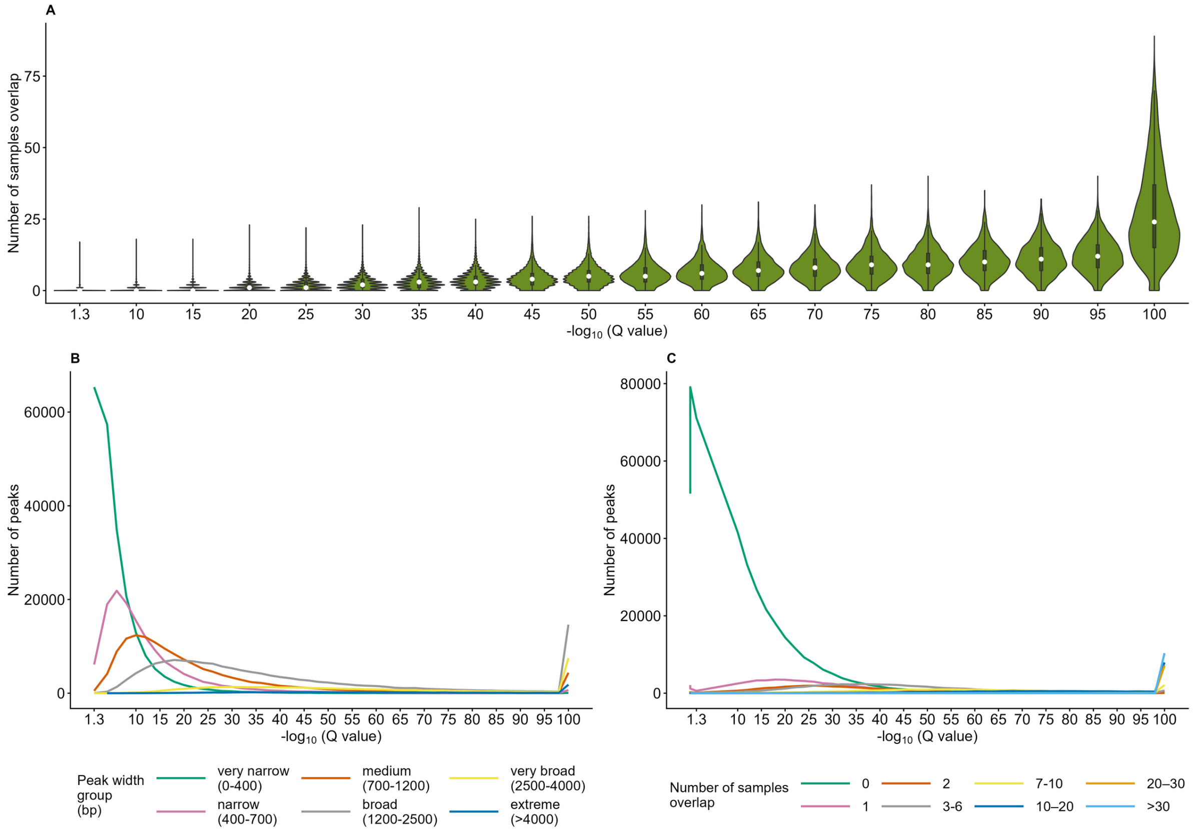

**Supplementary Figure 13: More stringent Q value thresholds alter the characteristics of detected peaks. A**) Violin plots showing the distribution of the number of samples overlapping each peak for peak sets generated at increasing Q value thresholds (x-axis). **B**) Line graph showing the distribution of peaks grouped by peak width across increasing Q-value thresholds. **C**) Line graph showing the distribution of peaks grouped according to the number of individual samples in which they are detected across increasing Q value thresholds. All peaks were called with Dataset 1 SOX10+ samples only.

### Supplementary Figure 14

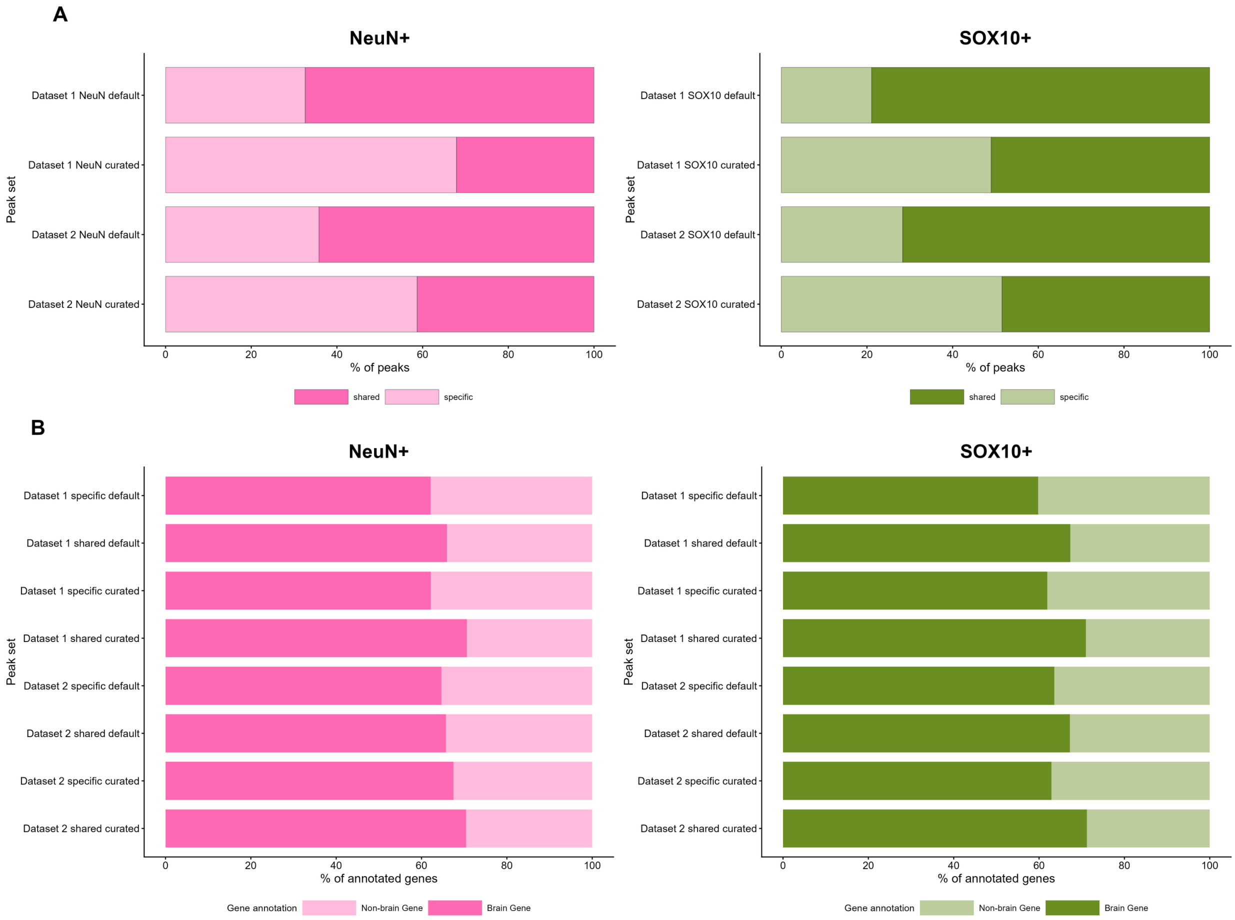

**Supplementary Figure 14: Curated peak sets are more reproducible across datasets and capture meaningful biological signal. A**) Stacked bar plots showing the percentage of peaks shared between datasets for the default peak sets ($Q < 0.05$) and curated peak sets ($Q <$ threshold determined by elbow analysis). Colours indicate whether peaks are shared between datasets or are dataset-specific, shown separately for each cell type. **B**) Stacked bar plots showing the percentage of genes in the default and curated peak sets that are classified as brain-expressed genes.
